# *In-vitro* cellular reprogramming to model gonad development and its disorders

**DOI:** 10.1101/2021.10.22.465384

**Authors:** Nitzan Gonen, Caroline Eozenou, Richard Mitter, Andreia Bernardo, Almira Chervova, Emmanuel Frachon, Pierre-Henri Commère, Inas Mazen, Samy Gobaa, Kenneth McElreavey, Robin Lovell-Badge, Anu Bashamboo

## Abstract

During embryonic development, mutually antagonistic signaling cascades determine the fate of the bipotential gonad towards a testicular or ovarian identity. Errors in this process result in human Disorders of Sex Development (DSDs), where there is discordance between chromosomal, gonadal, and anatomical sex. The absence of an appropriate, accessible *in-vitro* system is a major obstacle in understanding mechanisms of sex-determination/DSDs. Here, we describe protocols for differentiation of mouse and human pluripotent cells towards gonadal progenitors. Transcriptomic analysis reveals that the *in-vitro*-derived murine gonadal cells are equivalent to E11.5 *in-vivo* progenitors. Using similar conditions, Sertoli-like cells derived from 46,XY human induced pluripotent stem cells (hiPSCs) exhibit sustained expression of testis-specific genes, secrete AMH, migrate and form tubular structures. The cells derived from a 46,XY DSD female hiPSCs, carrying a *NR5A1* variant, show aberrant gene expression and absence of tubule formation. CRISPR/Cas9-mediated correction of the variant rescued the phenotype. This is a robust tool to understand mechanisms of sex-determination and model DSDs.

## Introduction

During embryonic gonad development the bipotential genital ridge adopts either a testicular or an ovarian cell fate. In the mouse, at around embryonic day (E) 10.0, somatic cells migrating from the coelomic epithelium overlaying the mesonephros, as well as from the latter, give rise to the bipotential gonad. Several key factors are expressed in the genital ridge at this stage including *Wt1* (Wilms’ tumor 1) (Kreidberg et al., 1993; Vidal and Schedl, 2000), *Sf1/Nr5a1* (Steroidogenic factor 1) (Luo et al., 1995; Luo et al., 1994), *Gatat4* (GATA Binding Protein 4) (Hu et al., 2013; Tevosian et al., 2002; Viger et al., 1998), *Cbx2* (Chromobox Protein homolog 2 / M33) (Garcia-Moreno et al., 2019b; Katoh-Fukui et al., 1998), *Fog2/ Zfpm2* (Friend of GATA protein 2) (Tevosian et al., 2002), *Lhx9* (Lim homeobox 9) (Birk et al., 2000) and *Emx2* (Empty Spiracles Homeobox 2) (Kusaka et al., 2010; Miyamoto et al., 1997). The developing gonad begins to become sexually dimorphic at the molecular level at around E10.7 (Garcia-Moreno et al., 2019a; Stevant et al., 2019; Stevant et al., 2018; Zhao et al., 2018) with the expression of *Sry* (Sex determining region on Y) in the supporting cell precursors, leading to their differentiation into Sertoli cells (Hacker et al., 1995; Koopman et al., 1990; Maatouk and Capel, 2008). Sertoli cells orchestrate testis cord formation and the differentiation of the other somatic testicular lineages as well as the primordial germ cells. SRY upregulates the expression of *Sox9*, a key transcription factor (TF), which, beyond a critical threshold, is both necessary and sufficient to induce and maintain Sertoli cells (Barrionuevo et al., 2006; Chaboissier et al., 2004; Foster et al., 1994; Gonen and Lovell-Badge, 2019; Gonen et al., 2017; Morais da Silva et al., 1996; Wagner et al., 1994). *Sox9*, along with other pro-testis factors (*Gata4, Wt1, Sf1, Dmrt1* and *Fgf9*) constitute a gene regulatory network that not only guides the precursor cells towards a testis fate, but also opposes the network required for the formation of ovarian cell types (Maatouk and Capel, 2008; Nef et al., 2019; Wilhelm et al., 2007).

Ovarian development requires *Rspo1/Wnt4/β-Catenin, Foxl2* and the recently identified *Runx1* signaling networks (Nicol et al., 2019). The RSPO1/WNT4 signaling pathway stabilizes β-catenin, which counteracts the establishment of a pro-testis *Sox9/Fgf9* network (Chassot et al., 2008; Kim et al., 2006; Maatouk et al., 2008; Vainio et al., 1999). In mice, FOXL2 is required to maintain ovarian identity postnatally, since the ablation of *Foxl2* in the murine adult ovary leads to trans-differentiation of granulosa cells into Sertoli cells (Uhlenhaut et al., 2009). *Runx1* has complementary/redundant roles with *Foxl2* to maintain fetal granulosa cell identity and combined loss of *Runx1* and *Foxl2* results in masculinization of fetal ovaries (Nicol et al., 2019).

Our understanding of sex-determination stems from embryological, genetic, and transcriptomic studies in mice and genetic analysis of humans with errors of sex determination. These individuals present with a phenotypic spectrum of conditions termed Disorders/differences of Sex Development (DSD), defined as congenital conditions in which development of chromosomal, gonadal, and anatomical sex is discordant (Bashamboo et al., 2017). However, a major bottleneck in further understanding mouse, but especially human sex-determination, is the lack of a robust *in-vitro* model that accurately recapitulates *in-vivo* gonad formation and development. Mouse models of variants associated with DSDs are limited not only by the expense and labour-intensive nature of producing mutant mouse lines (Bashamboo et al., 2017; Croft et al., 2017), but also by insufficient evolutionary conservation in sex-determination between mice and humans (Bashamboo et al., 2018; Miyado et al., 2016; Warr et al., 2016). Primary gonadal cells do not survive for long and their gene expression patterns rapidly diverge from normal in standard *in-vitro* culture conditions (Buganim et al., 2012). Although several Sertoli and granulosa cell lines have been established, these differ in expression profiles compared to their *in-vivo* counterparts, limiting their usefulness (Beverdam et al., 2003). Recently, several attempts have been made to generate Sertoli and Leydig cells *in-vitro* starting from pluripotent/multipotent cells, either by introducing exogenous transcription factors or using sequential supplement enriched media. Direct reprogramming of fibroblasts has also been attempted. However, the resultant populations lack appropriate and sustained expression of key testis-specific markers (Buganim et al., 2012; Knarston et al., 2020; Lan et al., 2013; Mae et al., 2013; Oeda et al., 2013; Rodriguez Gutierrez et al., 2018; Seol et al., 2018; Yoshino et al., 2021). No attempts were made to assess the entire transcription profiles, make comparisons to *in-vivo* counterparts, or, with the exception of Yoshino et al., (Yoshino et al., 2021) to determine the functional ability of the *in-vitro* derived gonadal cells. Moreover, none of these studies attempted to model DSDs.

Here, we developed a robust protocol for sequential differentiation of mouse embryonic stem cells (mESCs) towards gonadal progenitors using defined medium. Analysis of the transcriptome by RNA-Seq shows that these *in-vitro*-derived cells are highly similar to progenitors of E11.5 mouse gonads. Using the conditions optimized on mESCs, we derived Sertoli-like cells from human induced pluripotent stem cells (hiPSCs) derived from a healthy 46,XY male. The *in-vitro* differentiated population shows sustained expression of testis-specific genes, secrete AMH, migrate and spontaneously aggregate. Sorted cells can organize into 3D tubular structures on a specially designed microfluidic chip. For the first time, we also show that this protocol can be successfully used to model aspects of gonadal development with a naturally occurring human sex-reversing gene variant *in-vitro*.

## Materials and Methods

We used mouse ESCs and hiPSCs to derive somatic cells of the developing gonad, using defined supplement enriched medium. The details of cells lines and methodology are in supplementary appendix.

## Results

### Deriving TESCO-CFP; R26-rtTA XY mESCs

To enable differentiation of mESCs towards gonadal progenitors, we developed protocols that simulate the known pathway of gonad development (Figure 1A), whereby pluripotent cells differentiate first to become early mesoderm progenitors followed by further regional specification towards the mesoderm from which the gonads originate, which includes intermediate mesoderm (IM) and associated coelomic epithelium.

**Figure 1.**
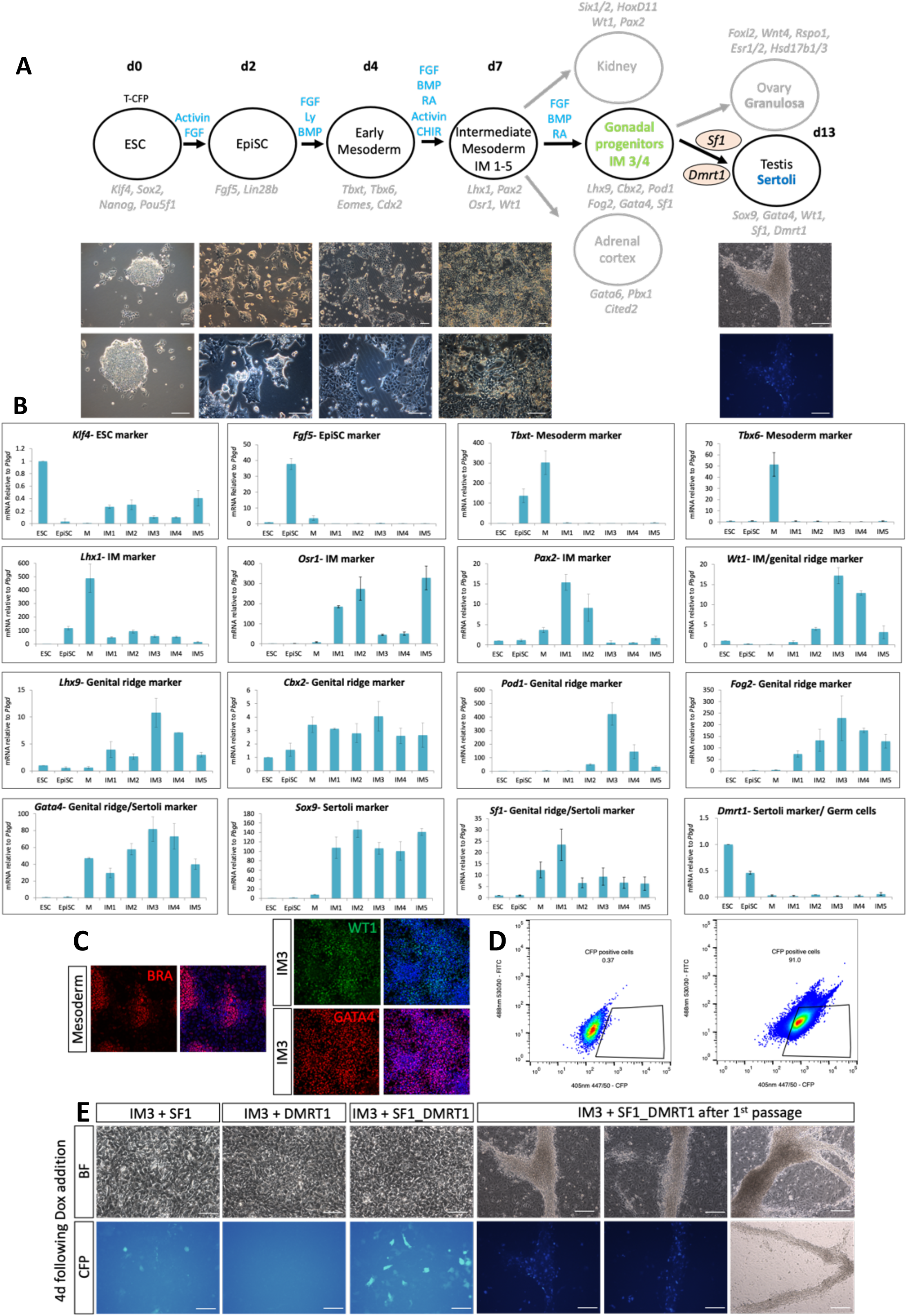
Differentiation of mouse ESCs towards gonadal cells. (A) Schematic representation of the differentiation protocol. T-CFP ESC (T-CFP; R26-rtTA, XY mESCs) were differentiated to EpiSCs followed by mesoderm induction, IM differentiation, and gonadal progenitor specification. Forced expression of *Sf1* and *Dmrt1* induced Sertoli-like cells expressing CFP which create tubule-like structures. Growth factors added at each step are highlighted in blue above the arrows. Markers of each stage are labelled in light grey. The timescale of the differentiation is presented at the top (day 0 - day 13). Representative bright field and fluorescent images are depicted below the schematic representation. Scale bars are 100 μm. (B) qPCR analysis of cells undergoing differentiation. Gene names are presented in the title. ESC, Embryonic stem cells; EpiSC, Epiblast stem cells; M, mesoderm; IM, Intermediate mesoderm. Data are presented as mean 2^-ΔΔCt^ values normalized to the house keeping gene *Pbgd*. Error bars show SD of 2^-ΔΔCt^ values. (C) Immunostaining of cells undergoing differentiation. Cells at the mesoderm stage were stained for BRACHYURY (Red). Cells at the IM3 stage were stained for GATA4 (Red) and WT1 (Green). Right hand images are merge with DAPI (blue). (D) Flow cytometry analysis showing CFP expression in IM3 differentiated cells (d13, left panel) and IM3 cells overexpressing *Sf1* and *Dmrt1* (d13, right panel). (E) Bright field and fluorescent images of IM3 cells overexpressing either *Sf1, Dmrt1* or both. Cells were imaged 4 days post Dox addition or after they were passaged (Ei), when tubule-like structures start to appear. Most CFP positive cells are located within tubules (Eii).

To facilitate the screening of culture conditions, we first derived ESCs from mice carrying the Sertoli cell-specific TESCO-CFP reporter (Sekido and Lovell-Badge, 2008). However, we also included the Rosa26-rtTA allele, which permits the forced expression, if required, of critical transcription factors using the doxycycline inducible system. Most experiments were performed using clone No. 7 which was XY, TESCO-CFP; R26-rtTA.

### Differentiating mESC towards gonadal progenitors

Differentiation of mESCs was initiated by passaging them in media containing 20% Knock Out Serum Replacement (KSR). Cells were then differentiated to an EpiSC-like state by culture in N2B27 media supplemented with 20ng/ml Activin, 12ng/ml bFGF and 1% KSR for 2.5 days as described by Hayashi et al. (Hayashi et al., 2011), who used a similar approach to generate Primordial Germ Cell-like cells (PGCLC) from mESCs. qPCR analysis of *Klf4* (Kruppel Like Factor 4), which is highly expressed in ESCs, but shows reduced expression in EpiSCs, and *Fgf5* (Fibroblast Growth Factor 5) which is induced in EpiSCs, confirmed that we are indeed able to generate EpiSCs (Figure 1B).

EpiSCs were then differentiated into early mesoderm progenitors, which express high levels of *Tbxt* (Brachyury, T-box Transcription Factor T) and *Tbx6* (T-box Transcription Factor 6), by exposing the cells to a chemically defined media supplemented with 20 ng/ml bFGF, 10 μM LY294002 and 10 ng/ml BMP4 (termed FLyB) for 36 hours, based on a protocol developed by Bernardo et al., (Bernardo et al., 2011). Robust mesodermal differentiation was evident by morphological changes in the cells (Figure 1A), qPCR analysis which showed high induction of *Tbxt/Bra* and *Tbx6* expression (Figure 1B), and immunostaining with an antibody against Brachyury/Tbxt, which showed that many of the cells are positive (Figure 1C). Relying on the BMP4 gradient to which cells in the region of the IM cells are exposed, we tested 5 different conditions (termed IM1-5) and screened the cells by marker analysis to identify the optimal protocol to obtain regional differentiation of the mesodermal cells to those most likely to be competent to form the gonads. All conditions used a defined media supplemented with bFGF (5 ng/ml) and medium levels of BMP4 (20 ng/ml). We then added additional factors including RA (100nM), Activin A (10 ng/ml) and CHIR99021 (WNT agonist, 0.3uM) (described in Materials and Methods). IM induction was performed for 3.5 days after which RNA was extracted and qPCR analysis performed for four major IM markers including *Lhx1* (Lim Homeobox 1), *Osr1* (Odd Skipped Related Transcription Factor 1), *Pax2* (Paired Box gene 2) and *Wt1* (Wilms’ tumor 1). All four factors were markedly induced, but to different extents (Figure 1B). While *Lhx1* was induced at the mesoderm stage and less so using the IM1-5 conditions, *Pax2* was strongly induced by both IM1 and IM2 conditions compared to the others. *Osr1* was induced by IM1, IM2 and IM5. *Wt1*, which is expressed in IM but continues to be expressed in the kidneys, supporting cell precursors of the gonad and later in Sertoli and granulosa cells, was mostly induced by IM3 and IM4. Since the IM markers show distinct expression profiles using the 5 differentiation conditions, we then determined the levels of early gonadal markers that are normally expressed at the bipotential stage of the gonads (E10.5-E11.5). These include *Lhx9, Cbx2, Pod1/ Tcf-21, Fog2/ Zfpm2* as well as *Gata4* (Wilhelm et al., 2007). All these factors were strongly induced in IM3 and to lesser extent in IM4 conditions, but at much lower levels in IM1, IM2 and IM5 conditions (Figure 1B). Immunostaining of IM3 cells with antibodies against WT1 and GATA4 indicated that most cells express high levels of these proteins (Figure 1C). This indicates that IM3 is the optimal condition to differentiate the mESCs towards early gonadal progenitors.

We also examined the expression levels of 3 additional TFs that are major markers of Sertoli cells and involved in Sertoli cell specification: *Sox9, Nr5a1* and *Dmrt1*. Surprisingly, all five IM conditions resulted in strong induction of *Sox9*, one of the most critical TFs of Sertoli cells, but one that is not unique to these (Figure 1B). *Nr5a1* and *Dmrt1* were not induced under any of the 5 conditions (Figure 1B). Although several Sertoli cell markers were naturally induced by the IM conditions, no CFP was apparent in these cells (Figure 1D, left panel), suggesting that Sertoli cell differentiation was incomplete.

### Transcriptomic analysis of the *in-vitro* derived early gonadal progenitors

To further verify the gene expression patterns of the various IM conditions in a more comprehensive and unbiased manner, we performed RNA-Seq analysis on the cells. Principal Component Analysis (PCA) showed the marked changes between ESC, EpiSC, M (Mesoderm) and the five IM conditions (IM1-5), which cluster closely together (Supplementary Figure 1A). Principal Component Analysis of only IM1-5 supported the results seen by the qPCR that IM3 and IM4 are quite similar, IM2 and IM5 are similar to each other while IM1 is distinct from both groups (Supplementary Figure 1B). The heatmap of the 100 most differentially expressed genes (50 top upregulated and 50 most downregulated) between IM1/2/5 and IM3/4 conditions included several interesting markers (Supplementary Figure 1C), such as the pluripotency marker *Sox2*, which is strongly expressed in ESC and EpiSC as expected. The three mesodermal markers *Tbxt*/*Bra* (*T*), *Cdx2* (*Caudal Type Homeobox 2*) and *Eomes* (*Eomesodermin*) were induced at the mesoderm stage. The IM/Sertoli marker *Wt1* was strongly expressed in IM3/4 and much less in IM1/2/5 (Supplementary Figure 1C) as confirmed by qPCR (Figure 1B). Lastly, the two metanephric mesenchyme markers, *Six2* (*SIX homeobox 2*) and *Hoxd11* (*Homeobox D11*) were more highly induced in IM2/5 compared to IM3/4 (Supplementary Figure 1C). We next generated a heatmap of known markers of the following lineages: ESC, EpiSC, mesoderm, paraxial mesoderm (PM), lateral plate mesoderm (LPM), IM, genital ridge, adrenal, kidney, Sertoli cells, granulosa cells and Leydig cells (gene list in Supp Table 3) and correlated expression of these markers with the various differentiation conditions (Supplementary Figure 1D). As expected, the pluripotency markers *Sox2, Nanog, Pou5f1/Oct4* and *Klf4* were highly expressed at the ESC and EpiSC stages. The EpiSC specific markers *Fgf5* and *Lin28b* were induced at the EpiSC stage. The mesodermal factors *Tbxt/Bra, Eomes, Tbx6, Cdx2* and *Mesp1* were strongly induced at the mesoderm (M) stage. Markers of the kidney and LPM lineages were induced in the IM1/IM2/IM5 conditions. In contrast markers of the genital ridge (e.g., *Wt1, Lhx9, Tcf21/Pod1, Cbx2, Gata4, Emx2* and *Zfpm2/Fog2*), and Sertoli cells (e.g., *Sox9, Fgf9, Erbb4, Cyp26b1, Cst9, Sox8* and *Shbg*) were highly induced in IM3 with slightly lesser induction at IM4 (Supplementary Figure 1D). Interestingly, transcripts from *Sry* were not detected in any of the conditions. *In vivo, Sry* shows a characteristic centre-to-pole wave of expression, and it is present very transiently in each cell (perhaps less than 3 or 4 hours) (Sekido et al., 2004); consequently, it may be missed in these *in vitro* conditions.

### Comparing *in-vitro*-derived gonadal progenitors with *in-vivo* gonadal cells

The combination of qPCR, immunostaining and RNA-Seq analysis strongly supports the hypothesis that IM3 differentiated cells correspond to early gonadal progenitors. To verify that this is indeed the case, we compared our transcriptomic data with that of mouse XY and XX gonads at E10.5, E11.5, E12.5 and E13.5 (Zhao et al., 2018). Comparing *in vivo-* versus *in-vitro*-derived cells, the immediate and major change at Principal Component 1 (PC1) was the separation of the *in-vitro* versus the *in-vivo* datasets (Supplementary Figure 2A). To overcome this limitation, we either examined the PC2/3 data (Supplementary Figure 2B) or used computational tools to correct the data (Figure 2A). The PC1/2 of the corrected data (Figure 2A) and the PC2/3 of the non-corrected data (Supplementary Figure 2B) indicated that in the IM condition cells cluster closely with early gonadal progenitors of E11.5 (Figure 2A, Supplementary Figure 2B).

**Figure 2.**
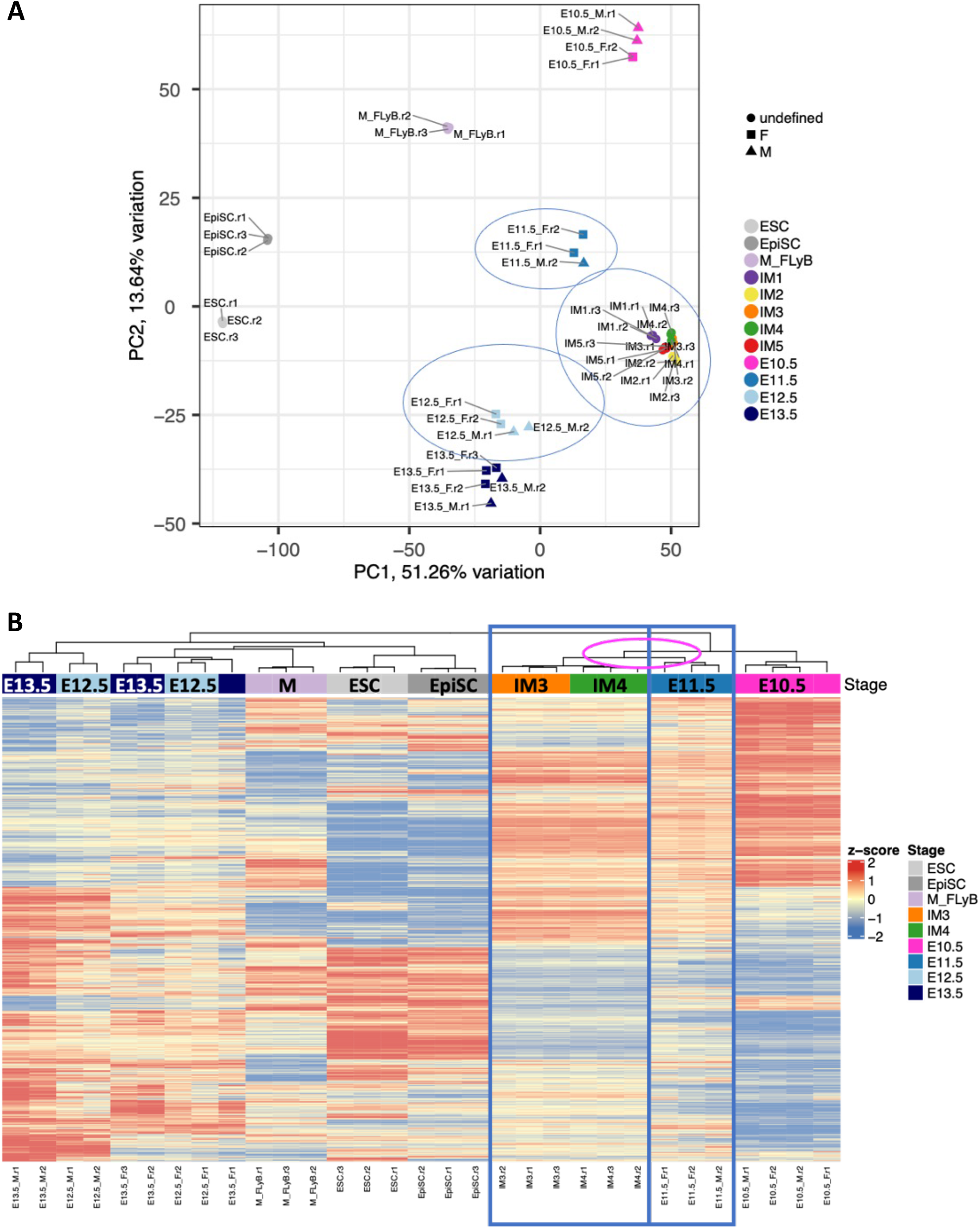
Comparing the transcriptome of *in vitro*-derived gonadal cells with *in vivo* gonadal cells. (A) Principal component analysis (PCA) of selected samples based on normalised mRNA expression level after batch correction. The top 2 PCs are shown. E10.5-E13.5 are gonadal cells isolated from entire embryonic male (triangle) or female (square) gonads (Zhao et al., 2018). Three biological replicates were analysed for each sample type. (B) Heatmap of the genes most differentially expressed between the E10.5 and E13.5 XY male gonads following batch correction (Zhao et al., 2018). IM3/IM4 cluster closely to the E11.5 *in vivo* gonadal cells. Gene-level expression across samples is shown as a z-score running from red (high) to blue (low). Three biological replicates were analysed for each sample type.

Using the Zhao et al. dataset we chose the genes most differentially expressed (DE) between E10.5 to E13.5 XY gonads, because these are the genes induced during gonadal differentiation. Analysing these genes with the different *in-vivo* and *in-vitro*-derived cells demonstrated that E11.5 bipotential gonads cluster with IM3 and IM4 cells and exhibit similar gene expression patterns (Figure 2B). The Poisson distance heatmap based on DE genes between E10.5 to E13.5 XY gonads shows a similar trend with IM3/4 being the most similar to early gonads (Supplementary Figure 2C). Selecting the most DE genes between E10.5 to E13.5 XX gonads, similar results were obtained both for Poisson distance heatmap (Supplementary Figure 2D) and for differential gene expression analysis (Supplementary Figure 2E). Altogether this data suggests that the IM3/IM4 conditions generate cells that are similar to E11.5 gonadal progenitors of the XY and XX gonads. This can serve as an optimal platform to generate all the somatic cells of the gonad.

### Inducing embryonic Sertoli cells from *in-vitro*-derived gonadal progenitors

We next examined whether we can further differentiate the early gonadal progenitors towards embryonic Sertoli cells. Forced expression of *Sox9, Wt1, Gata4, Sf1* and *Dmrt1* was shown to induce fibroblasts to differentiate into embryonic Sertoli-like cells (Buganim et al., 2012). This, along with other studies, suggests that these five TFs are the critical players for Sertoli cell fate (Wilhelm et al., 2007). Three out of the five TFs (*Gata4, Wt1* and *Sox9*) were naturally induced in our differentiation protocol using growth factors, yet *Sf1* and *Dmrt1* were not induced. The Sertoli cell-specific reporter TESCO-CFP was also not activated in the cells using the IM conditions. Therefore, we examined whether forced overexpression of *Sf1* and *Dmrt1* can induce embryonic Sertoli cell differentiation from the early gonad progenitors. To that aim we used doxycycline-inducible expression vectors and generated lentiviruses which overexpress either factor. IM3 cells were infected with either *Sf1*/*Dmrt1*-expressing lentiviruses or a combination of both and Doxycycline was added for 4 days (Figure 1E). Overexpression of *Dmrt1* alone did not lead to expression of the TESCO-CFP reporter while overexpression of *Sf1* alone resulted in weak CFP signals in several cells (Figure 1E). On the contrary, overexpression of both *Dmrt1* and *Sf1* resulted in strong expression of the TESCO-CFP reporter (Figure 1E). Analysing the cells using flow cytometry indicated that, compared to control IM3 cells (without overexpression of the two factors), ~90% of IM3 cells with forced expression of *Sf1* and *Dmrt1* were CFP positive (Figure 1D, right panel). Upon further passaging of the cells, they started forming tubule-like structures, reminiscent of testis cords. Remarkably, many of the tubule forming cells were CFP positive, and are therefore likely to resemble Sertoli cells (Figure 1A, 1E).

To determine how similar the CFP positive cells were to embryonic Sertoli cells, we sorted them by flow cytometry and isolated RNA, which was analysed by RNA-Seq. Because the Zhao et al., (Zhao et al., 2018) dataset used entire XY and XX gonads, rather than purified Sertoli cells, this was not a very relevant comparison. We therefore used the dataset from Maatouk et al. (Maatouk et al., 2017), which has RNA-Seq data from TESCO-CFP sorted E15.5 Sertoli cells. We generated a list of DE genes between E13.5 XY and XX gonads as this should indicate Sertoli versus granulosa cell markers. The heatmap of these DE transcripts indicated that the induced Sertoli-like cells (iSLCs) cluster next to the E15.5 Sertoli cells (SC), but the datasets are not overlapping (Figure 3A). This suggests that the iSLCs resemble fetal Sertoli cells, but that they are not identical to the latter. A Poisson distance heatmap generated from the same transcripts indicates that the iSLCs are more similar to E12.5 and E13.5 male gonads (Figure 3B, black arrows) than to E12.5 and E13.5 female gonads (Figure 3B, white arrows).

**Figure 3.**
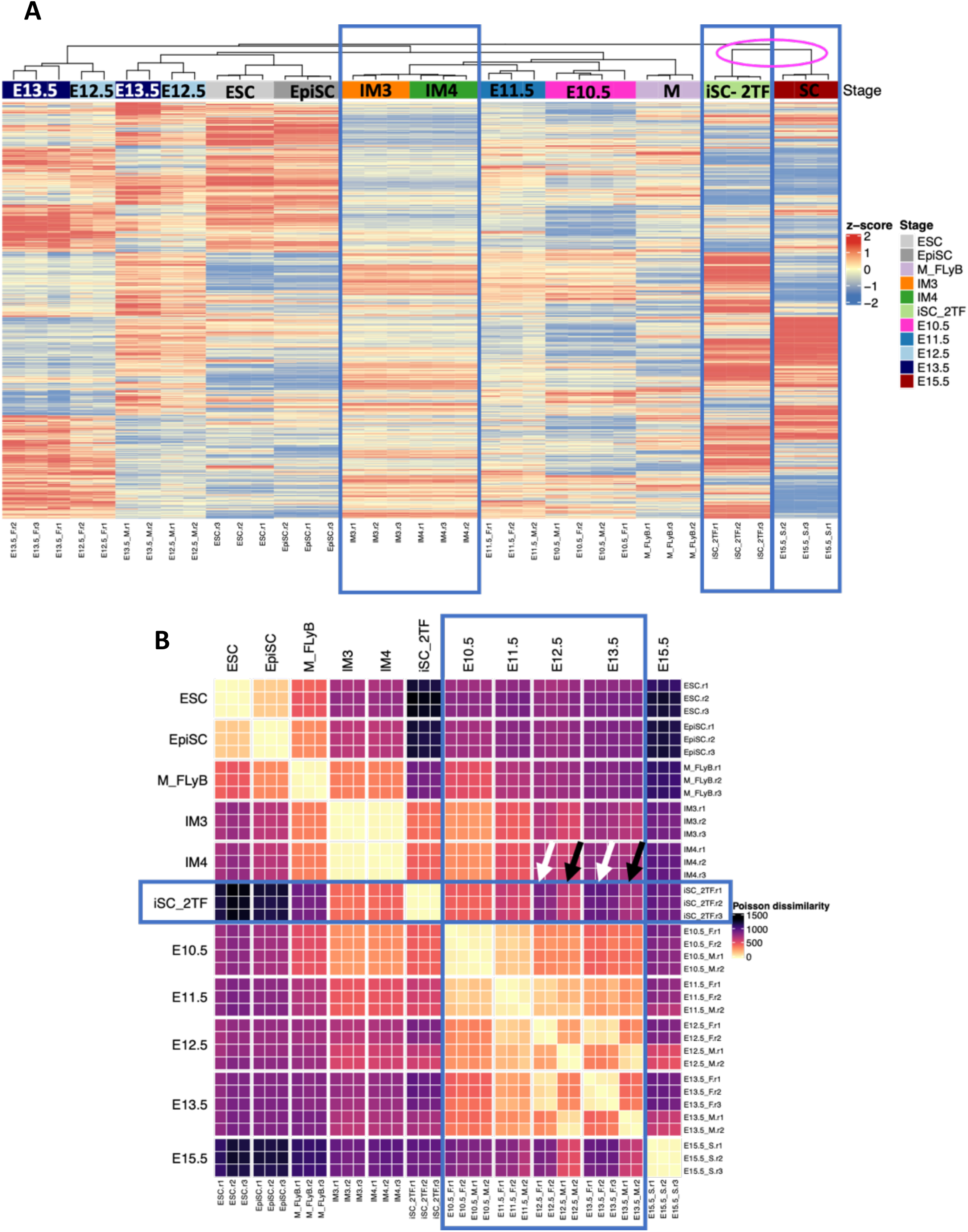
Comparing the transcriptome of *in vitro*-derived gonadal cells with *in vivo* gonadal cells and E15.5 purified Sertoli cells. (A) Heatmap of normalised abundance after batch correction for genes differentially expressed between E13.5 XY and E13.5 XX gonads. Induced Sertoli-like cells following forced expression of *Sf1* and *Dmrt1* (iSC-2TF) cluster closely to the E15.5 TESCO-CFP (Sekido and Lovell-Badge, 2008) purified Sertoli cells (Maatouk et al., 2017), however the gene expression pattern indicates that the two are not identical. (B) Heatmap of Poisson dissimilarity scores showing transcriptional similarities between samples. Dissimilarity scores were calculated from a normalised abundance matrix of genes differentially expressed between E13.5 XY and E13.5 XX gonads (Zhao et al., 2018). Blue boxes indicate the similarity between induced Sertoli-like cells (iSC-2TF) and the E10.5-E13.5 *in vivo* gonadal cells. The iSC-2TF are more similar to male-derived gonadal cells (black arrows) than to female derived gonadal cells (white arrows). Relative sample dissimilarity is shown running from dark purple (dissimilar) to yellow (similar).

Altogether this data represents the first approach at generating early gonadal progenitors that have then been compared to cells from *in vivo*-derived gonads. The data shows that the two are highly similar and hence the *in vitro*-derived cells constitute excellent starting material for further differentiation of different somatic gonadal cell types. Our induced Sertoli-like cells activate the TESCO-CFP reporter and can form tubule like structures, reminiscent of Sertoli cells.

### Deriving supporting cells of the gonad from hiPSCs derived from 46,XY male, 46,XX female and a 46, XY DSD patient

We next determined if it was possible to use human induced Sertoli-like cells (hiSLCs) in these protocols in order to model a pathogenic variant which causes male-to-female sex-reversal. To differentiate the hiSLCs we used the protocol optimized on mouse ES cells with minor modifications to the medium composition. We used three different hiPSC lines (2 clones each): two from healthy individuals (46,XY male and 46,XX female) and one from a patient with 46,XY DSD that carries the pathogenic p.Arg313Cys variant in *NR5A1*. Heterozygous pathogenic variants in *NR5A1* are one of the most common causes of 46,XY DSD. The phenotypes associated with *NR5A1* are highly variable and range from a complete absence of testis-determination (46,XY gonadal dysgenesis) and female external genitalia to testes in individuals with typical male external genitalia, but who are infertile.

During the differentiation process, cells exhibit changes in shape and transcript levels (Figure 4A and B). After 12 days of sequential culture in defined medium, cells from the control XY male show rearrangement and aggregation in the dish (Figure 4A), which is not visible for the two other lines.

**Figure 4.**
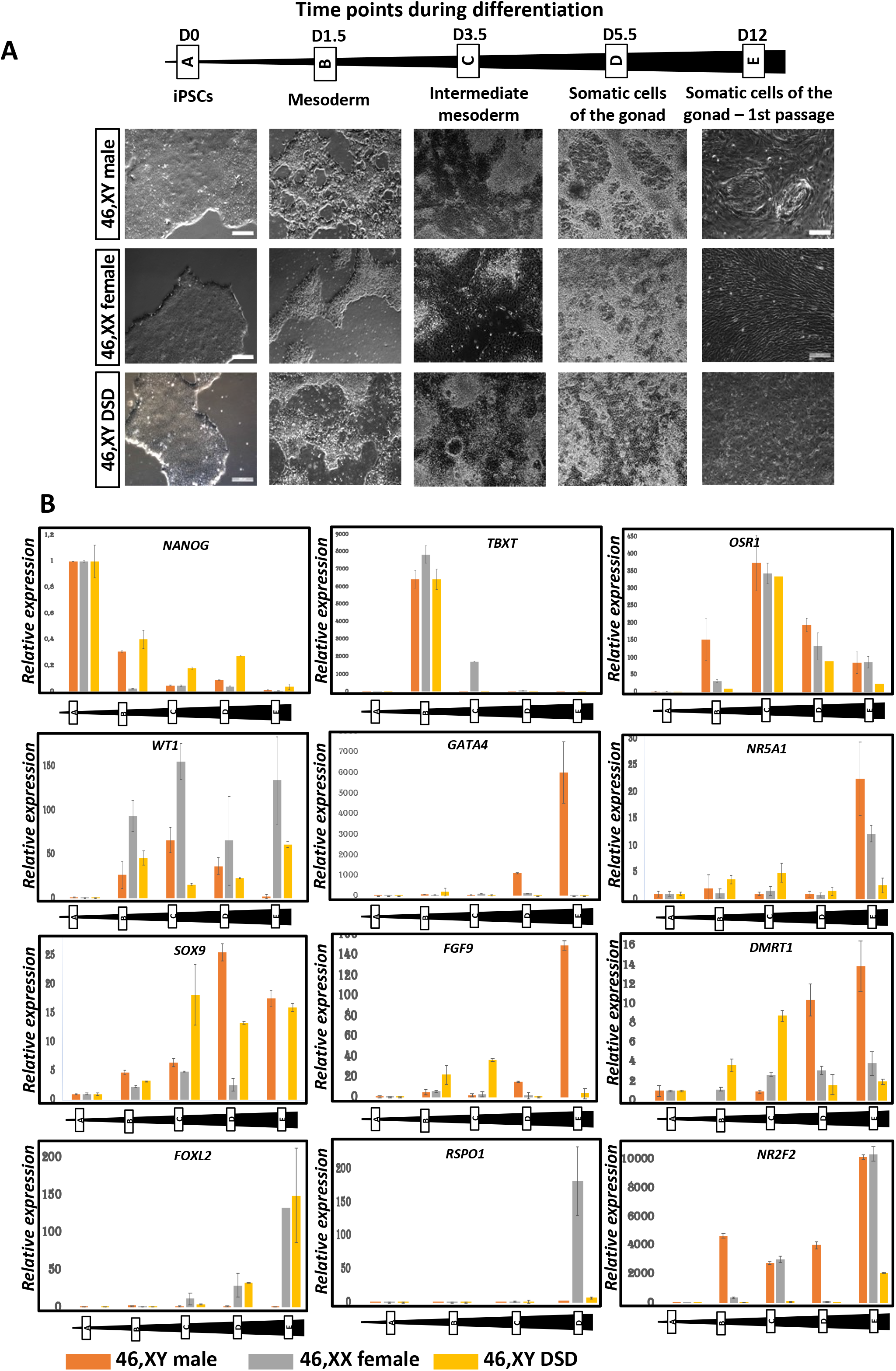
Differentiation of human iPSCs derived from healthy 46,XY male, 46,XX female and a 46,XY DSD patient with a pathogenic variant in *NR5A1*. (A) Differentiation time points are indicated together with BF images of the cells at each stage of the differentiation process. iPSC derived from the 46,XY healthy male are the only cells to self-aggregate and form tubule-like structures. D, Days, Solid bar. (B) Representative RT-qPCR expression profiles of selected differentiation markers of the three iPSC-derived cell lines during the differentiation process.

RT qPCR of the three cell lines (Figure 4B) reveals that expression of the pluripotency marker *NANOG* gradually reduces during the differentiation process. The cells differentiate into mesoderm and then IM as demonstrated by the expression of *T* (*TBXT*/*BRA*) and *OSR*1 respectively (Figure 4B). *WT1* expression showed a gradual reduction in the 46,XY derived cell cultures, but persisted in those derived from 46,XX and 46,XY DSD cells (Figure 4B). As differentiation continues, the somatic cells derived from 46,XY hiPSCs express markers specific to the somatic cells of the male gonad (*GATA4, NR5A1, DMRT1, SOX9* and *FGF9*,) whereas the 46,XX cells express markers specific for the somatic cells of the female gonad (*FOXL2, RSPO1*) (Figure 4B, Supplementary Figures 3A,B,C). The somatic cells derived from a 46,XY DSD patient carrying the *NR5A1* variant p.Arg313Cys, show aberrant expression of gonadal transcripts with either a reduction (*NR5A1, DMRT1*) or an absence (*GATA4, FGF9*) of expression of Sertoli markers and at the same time there is increased expression of the granulosa marker *FOXL2* (Figure 4B). Interestingly, the cells derived from 46,XY DSD hiPSCs fail to express *FGF9* even though *SOX9* is expressed, suggesting a breakdown in establishment of the SOX9/FGF9 feed forward loop required for the maintenance of Sertoli cell identity. The somatic cells derived from both 46,XX and 46,XY iPSC express *NR2F2*, a marker of presumptive steroidogenic cells, which is considerably reduced in the cells derived from 46,XY DSD hiPSCs (Figure 4B).

### Functional and structural characterization of the somatic cells derived from hiPSCs

AMH, which is required for the regression of the Müllerian duct derivatives (Josso, 1992), is secreted by fetal Sertoli, but not granulosa cells. Consistent with this, somatic cells derived from 46,XY iPSCs secrete AMH (Figure 5A). On prolonged culture (for over 7 weeks), cells derived from both 46,XY and 46,XY DSD hiPSCs continued to express the SOX9 protein (Figure 5B. Supplementary Figure 4) and other Sertoli cell markers (NR5A1, DMRT1; Supplementary Figure 4). Following 12-15 days of sequential culture in defined media, the cells derived from 46,XY hiPSCs spontaneously form tubular structures (Figure 5C, top panel). These tubules are composed of SOX9 positive cells (Figure 5C, lower panels), further suggesting that they are Sertoli-like. In the developing testis CLAUDIN-11 is a critical transmembrane component of Sertoli cell tight junctions and is imperative for initiation and maintenance of functional Sertoli cell differentiation (Morita et al., 1999). Murine Sertoli cells lacking *Claudin-11* can proliferate and maintain the expression of testis cell markers, however, they acquire a fibroblast phenotype, lose tight junction integrity, and are eliminated through the lumen (Gow et al., 1999). The Sertoli-like cells derived from 46,XY and 46,XY DSD hiPSCs express both SOX9 and CLAUDIN-11 (Figure 5D). However unlike the cells derived from 46,XY hiPSCs, those derived from 46,XY DSD hiPSCs lose the organized spatial distribution of CLAUDIN-11 surrounding the SOX9 expressing cells. This leads to the disruption of tight junctions resulting in absence of tubule formation (see below). The somatic cells derived from 46,XY hiPSCs express SOX9 and those from the 46,XX hiPSCs express FOXL2. In contrast to the cells derived from control individuals, those from the 46,XY DSD iPSCs express both the pro-testis SOX9 and pro-ovary FOXL2 proteins (Figure 5E). Similar cell populations were identified in adult murine testes after *Dmrt1* ablation (Matson et al., 2011). These cells possibly represent multipotent progenitors or cells transitioning from Sertoli cells to granulosa cells in the trans-differentiating gonad.

**Figure 5.**
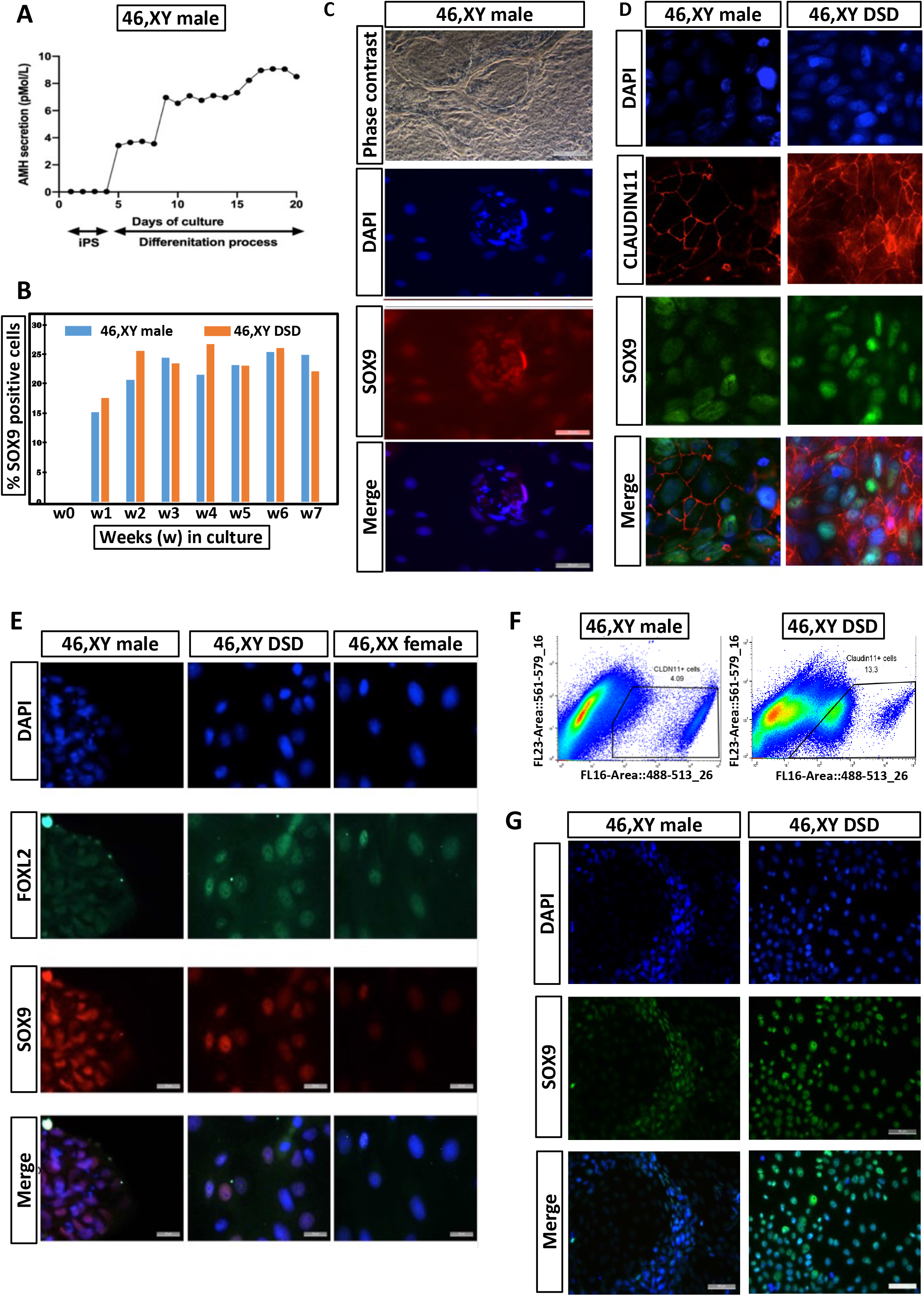
Characterisation of iSLCs from 46,XY male and 46,XY DSD patient. (A) The Sertoli cell product, AMH, is secreted into the medium by the iPSC-derived 46,XY male cells during differentiation. (B) Continuous and prolonged expression of SOX9 in the differentiating 46,XY male and 46,XY DSD cells. (C) Top panel: phase contrast of iPSC-derived 46,XY cells after 12 days of sequential culture in conditioned media, shows spontaneous formation of tubular structures. Middle and bottom panels: the tubule-like structures are composed of SOX9 positive cells. (D) Expression of SOX9 and CLAUDIN11 in iPSC-derived cells from 46,XY healthy male and 46,XY DSD patient after 12 days of sequential culture in defined media. Both cell lines express SOX9 and CLAUDIN11, but the spatial organisation of CLAUDIN11 is disrupted in iPSC-derived cells from the 46,XY DSD patient. (E) Immunocytochemistry for SOX9 and FOXL2 in iPSC-derived cells from 46,XY healthy male, 46,XX female and 46,XY DSD patient after 12 (+5) days of sequential culture in defined media. The iPSC-derived 46,XY male cells express SOX9 but not FOXL2, whereas the iPSC-derived 46,XX cells express FOXL2 but not SOX9. In contrast, a proportion of the iPSC-derived 46,XY DSD cells express both SOX9 and FOXL2 within the same cell. (F) Flow cytometric sorted iPSC-derived CLAUDIN11-positive cells from a 46,XY male and 46,XY DSD patient following 12 (+12) days of sequential culture in defined media. (G) After FACS sorting, the iPSC-derived CLAUDIN11-positive cells from a 46,XY male and a 46,XY DSD patient are seeded onto Matrigel in a chamber slide. After 7-10 days, the 46,XY male cells spontaneously organise into circular tubules whereas 46,XY DSD cells do not form these structures. Both the cell lines express SOX9.

During the differentiation process from 46,XY and 46,XY DSD hiPSCs, CLAUDIN-11 is expressed in SOX9 positive cells (Figure 5D). We sorted CLAUDIN-11 positive cells by flow cytometry following 12 (+12) days of sequential culture in supplement enriched media (Figure 5F). The sorted cells express SOX9 and were seeded on Matrigel coated chamber slides. After 7-10 days in culture the 46,XY cells spontaneously organize into circular tubules whereas 46,XY DSD cells are unable to form organised structures (Figure 5G), possibly due to disruption in organisation of CLAUDIN-11.

To further assess the ability of hiPSC-derived supporting cells to form tubules we seeded the sorted cells (after expansion for 2-3 weeks) atop a soft substrate made of Matrigel (50% v/v). The cells derived from 46,XY iPSCs can form tubules whereas those derived from 46,XX and 46, XY-DSD hiPSCs aggregate but fail to form tubules (Figure 6A). To evaluate the capacity of supporting cells derived from iPSCs to migrate, aggregate and form 3D structures, we designed a poly-dimethyl-siloxane (PDMS) microfluidic device termed GONAChip (Figure 6B) with three channels: one central, for the Matrigel loading, and two lateral channels for media and cell loading respectively (Figure 6C). Gaps in the channel walls provide a medium/Matrigel interface on which cells where cultured. After sorting, the CLAUDIN-11 positive cells from 46,XY, 46,XX and 46,XY DSD lines were expanded for 2-3 weeks. Cells were then seeded on one lateral channel of GONAChip. The seeded devices where then incubated in a humid chamber at 37°C in 5% CO_2_ for 72 hours. Live imaging (Supplementary videos 1,2 and 3) shows that cells from all the three cell lines can survive and proliferate. We found that 46,XY and 46,XY DSD cells migrate and invade the central channel filled with Matrigel (hours 0-30) whereas the 46,XX cells do not. We also found that 46,XY cells seem to migrate in a coordinated manner that leads to recognizable tubular structures. On the other hand 46,XY DSD invasion of Matrigel seemed less coordinated (Hours 30-72) and the cells do not form tubules (Figure 6D).

**Figure 6.**
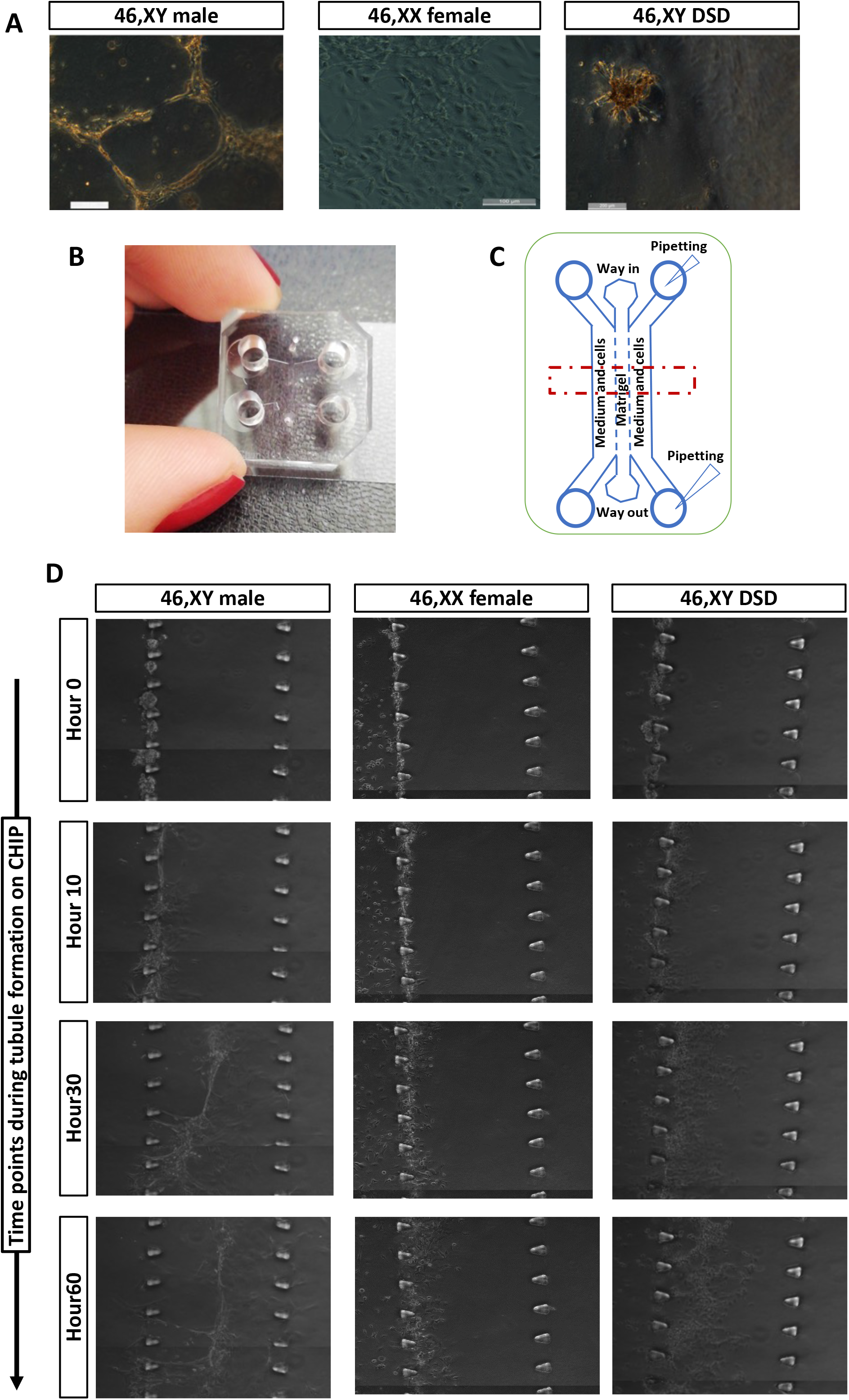
3D tubule formation in a microfluidic chip. (A) After FACS sorting, the iPSC-derived CLAUDIN11-positive cells from 46,XY male, 46,XX female and 46,XY DSD cells were expanded for 7-14 days in Sertoli cell medium. These cells were then seeded on a constrained space on hardened (50%) Matrigel domes. The iPSC-derived cells from the 46 XY male migrate and form tubules whereas, 46,XX female and 46,XY DSD cells do not form tubular structures. (B) Poly-dimethyl-siloxane (PDMS) microfluidic Chip designed for 3D tubule formation (GONAChip) together with, (C) A schematic representation of the Chip. (D) Still images from time laps video of cell migration, aggregation, and tubule formation on GONAChip. The time lapse videos (Supplementary videos 1,2 and 3) show the cells from all the three iPSC-derived cell lines can proliferate. The iPSC-derived 46,XX cells do not migrate whereas 46,XY male and 46,XY DSD cells migrate and enter the central chamber lined with Matrigel (hour 0-30). The iPSC-derived 46,XY male cells organise and form distinct tubular structures, whereas the iPSC-derived 46,XY DSD cells do not form tubular structures (Hour 30-72).

### CRISPR/Cas9 genome-edited 46,XY DSD cells resemble 46,XY cells

We corrected the pathogenic variant in 46,XY DSD hiPSCs using CRISPR/Cas9 genomeediting. The edited cells were then subjected to differentiation following the same protocols as for the 46,XY and 46,XY DSD iPSCs (Figure 7A). The pluripotent cells (expressing *NANOG*) differentiate into mesoderm (expressing *T*) and intermediate mesoderm (expressing *WT1*). As differentiation occurs these cells express markers specific to the somatic cells of the male gonad (*NR5A1, DMRT1, SOX9*) (Figure 7B, 7C). Interestingly the 46,XY DSD cells after correction of the *NR5A1* variant, regain the expression of *FGF9* while *FOXL2* expression cannot be detected (Figure 7B, 7C). Moreover, the cells show spatial reorganisation of CLAUDIN-11 similar to that of 46,XY cells (Figure 7C). On prolonged culture (for over 7 weeks) the cells derived from this rescued cell line continue to express the SOX9 protein (Figure 7D), NR5A1 and DMRT1 (Supplementary Figure 4).

**Figure 7.**
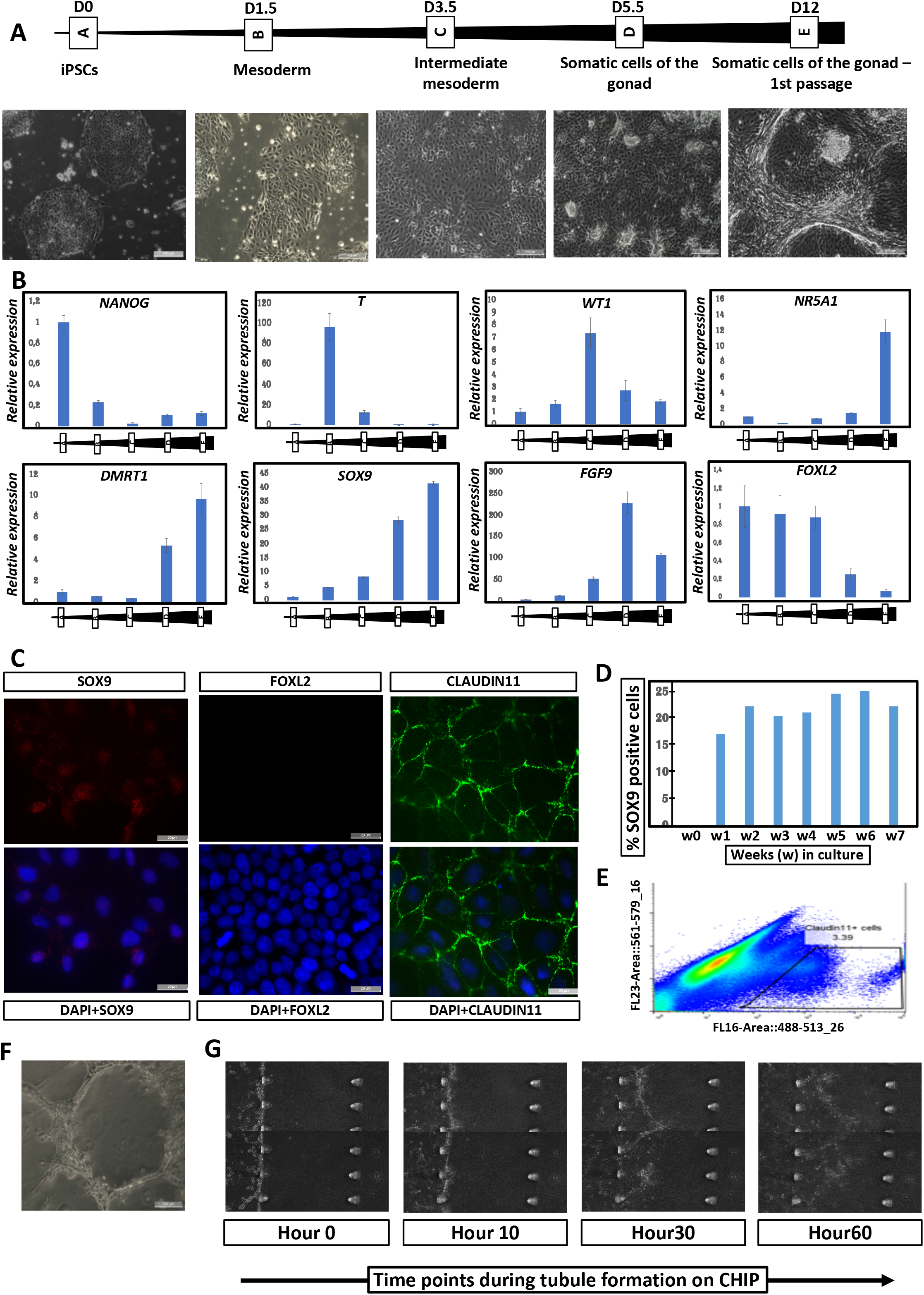
CRISPR/CAS9 correction of the *NR5A1* pathogenic variant in iPSC-derived 46,XY DSD cells restores iSLC properties. (A) Differentiation time points and BF images of the corrected 46,XY DSD cells during the differentiation process. (B) RT qPCR expression data of selected markers during the differentiation process. (C) Expression of SOX9, FOXL2 and CLAUDIN11 in *NR5A1*-corrected 46,XY DSD iPSC-derived cells after 12 days of sequential culture in defined media. Cells express SOX9 and CLAUDIN11, but the expression of FOXL2 is not detected. Moreover, CLAUDIN11 expression is spatially organised, similar to that seen in 46 XY, male cells (Figure 5D). (D) Continuous and prolonged expression of SOX9 is observed in the differentiating cells *NR5A1*-corrected 46,XY DSD iPSC-derived cells. (E) Flow cytometric sorting of CLAUDIN11 positive *NR5A1*-corrected 46,XY DSD iPSC-derived cells following 12 (+12) days of sequential culture in conditioned media. (F) After FACS sorting, the CLAUDIN11 positive *NR5A1*-corrected 46,XY DSD iPSC-derived cells were expanded for 7-14 days in Sertoli cell medium, then seeded on a constrained space on hardened (50%) Matrigel domes. The cells migrate and form tubules. (G) Still images from a time lapse video of cell migration, aggregation, and tubule formation on GONAChip (Supplementary video 4) show that the cells proliferate, migrate (Hour 0-30), and self-organise to form distinct 3D tubular structures (Hour 30-72).

The rescued cells were sorted using CLAUDIN-11 (Figure 7E), expanded, and seeded atop Matrigel substrates as described above. These cells show spontaneous tubule formation similar to those derived from 46,XY hiPSCs (Figure 7F). Using the GONAChip device, sorted CLAUDIN-11-positive rescued cells proliferated, migrated (Hour 0-30) and self-organized to form distinct 3D tubular structures (Hour 30-72) similar to those derived from 46,XY DSD iPSCs (Images extracted from time lapse videos Supplementary video 4).

## Discussion

Several attempts have been made for *in-vitro* generation of embryonic gonadal somatic cells either by forced expression of exogenous transcription factors (Buganim et al., 2012) or using combinations of growth factors and defined media (Mae et al., 2013; Oeda et al., 2013; Seol et al., 2018). In most of these studies, the resulting populations were defined as Sertoli cells based on expression of certain Sertoli-specific markers. However, they show a limited activation of endogenous Sertoli cell markers, with little to no activation of the key testis factor *Sox9*. An attractive route is the use of supplement enriched media to differentiate human pluripotent cells towards the IM and associated coelomic epithelium (Takasato and Little, 2016) and further towards granulosa-like (Lan et al., 2013) or Sertoli-like cells (Knarston et al., 2020; Rodriguez Gutierrez et al., 2018). However, the *in-vitro* derived Sertoli-like cells also lack robust and prolonged expression of key Sertoli-cell markers (SRY, SOX9, NR5A1, DMRT1) and their functionality was not unexplored. This precluded their use as a sufficiently accurate model to study naturally occurring variants causing errors in human testis-determination.

We have derived gonadal progenitors from mouse ESCs and human iPSCs following the *in-vivo* developmental pathway for gonad formation. For differentiation of mESCs to IM, we optimized the culture conditions where medium contained bFGF, BMP and RA but not the WNT agonist CHIR99021 or Activin A. The IM cells were further differentiated towards the somatic cells of the gonad. In contrast to other studies, the transcriptomic profile of our *in-vitro*-derived mouse gonadal cells strongly resembled that of E10.5-E11.5 gonadal progenitors (Zhao et al., 2018). The expression of three key TFs (*Gata4, Wt1* and *Sox9*) was endogenously induced in the progenitor cells, whereas that of *Nr5a1* and *Dmrt1* was not. Forced overexpression of *Nr5a1* and *Dmrt1* in the *in-vitro*-derived progenitors resulted in strong activation of the Sertoli cell-specific TESCO-CFP reporter (Sekido and Lovell-Badge, 2008) and formation of tubule-like structures resembling testis-cords. The transcriptome of the CFP-positive *in-vitro*-derived progenitor cells was similar to *in-vivo* TESCO-CFP E15.5 Sertoli cells, however these were not identical. This could be because our *in-vitro*-derived Sertoli-like cells are less mature than E15.5 Sertoli cells and comparisons with earlier stages, such as E12.5 or E13.5, may show higher degrees of similarity. This opens an avenue for future research to explore conditions that promote maturation of Sertoli cells.

Using the culture conditions optimised with mESCs, we directed both 46,XY and 46,XX hiPSCs towards gonadal progenitors. Although similar growth factors were used, we observed differences in gene expression not only between the mouse and human derived cells but also between our hiPSCs-derived cells and other published studies. Notably, *Sry* transcripts were absent in all conditions tested using the murine cells although both *SRY* mRNA and protein were detected at low levels in XY hiPSCs and in the human XY *in-vitro*-derived gonadal cells (Supplementary Figure 5) respectively. This also contrasts with the study by Knarson et al. (2020) where *SRY* mRNA expression was not detected in hiPSCs-derived Sertoli-like cells. SOX9 overexpression can drive testis differentiation in the absence of *Sry* in transgenic XX embryos (Vidal et al., 2001) or when the *SOX9* gene or *SOX9*-specific regulatory elements are duplicated (Croft et al., 2018; Huang et al., 1999; Qian et al., 2021). It is possible that the supplement enriched medium used by us for differentiation of mESCs and by others for differentiation of hiPSCs, bypasses *SRY* expression and directly or indirectly upregulates *SOX9*. However, we observed that cells derived from 46,XX hiPSC lines did not express Sertoli cell markers. This suggests that derivation of Sertoli-like cells from hiPSCs in our culture system is dependent on SRY as would usually occur *in vivo*. We also observed differences in the expression of two other key testis-determining genes *NR5A1* and *DMRT1*. These genes were not induced in the gonadal progenitors derived from mESCs. Similarly, in the study by Knarson et. al. (2020), *NR5A1* and *DMRT1* expression was either not induced or expressed at very low levels. This could be because induction of *DMRT1* expression is contingent on a threshold of SRY expression. In contrast to previous studies, we observe a robust and prolonged (for over 7 weeks) expression of the key testis-determining genes, *SOX9*, *NR5A1* and *DMRT1* in hiPSC-derived Sertoli-like cells. In monolayer cultures, these cells secrete AMH, self-aggregate and spontaneously form tubule-like structures reminiscent of testis cords. The tubular structures are lined with SOX9-positive cells surrounded by the tight junction protein CLAUDIN-11. Using a specially designed poly-dimethyl-siloxane (PDMS) microfluidic chip we demonstrated that CLAUDIN11-sorted *in-vitro*-derived Sertoli-like cells can migrate and self-aggregate to form 3D tubules.

We can direct pluripotent cells from both mouse and human, to mimic the *in-vivo* developmental pathway, to generate early gonadal progenitors and Sertoli-like cells using sequential defined media. This approach circumvents the need for continuous expression of exogenous factors involved in sex-determination (Buganim et al., 2012; Liang et al., 2019; Xu et al., 2020), which can override the endogenous signals and force these cells to develop into the somatic cells of the gonad. By allowing the cells to differentiate, based on their natural genetic composition, enables the investigation of naturally occurring gene variants of DSD that disrupt this process (Bashamboo et al., 2017).

As a proof of principle, we differentiated iPSCs from a DSD patient carrying a naturally occurring sex-reversing missense variant. The somatic cells derived from the patient’s iPSCs did not show appropriate stage-specific expression of Sertoli cell markers and lacked the ability to self-aggregate or form testis tubule-like structures. Correction of this missense variant by CRISPR/Cas9 in patient’s iPSCs was sufficient to re-establish the ability of the derived cells to form Sertoli-like cells that can aggregate to form tubules similar to the cells derived from 46,XY male.

This *in vitro* system can therefore be used to study human testis-determination as well as the effects of naturally occurring variants that disrupt this processes and cause DSD. This is important as >50% of all DSD cases do not have an established genetic etiology and the phenotypic variability associated with variants in known DSD genes is currently unexplained (Bashamboo et al., 2017). This would enable us to determine both the causality of novel genes proposed to cause DSD, and explore the phenotypic variability observed between individuals with identical pathogenic gene variants.

We can generate high quality early gonadal progenitors and Sertoli-like cells. The system could be refined further to produce other somatic lineages of the gonad, such as Leydig and peritubular myoid cells of the testis or granulosa and theca cells of the ovary, as well as other gonadal lineages that are being identified by single-cell sequencing approaches (Mayere et al., 2021; Stevant et al., 2019; Stevant et al., 2018). Recently, foetal ovarian somatic cell-like cells (FOSLCs) were derived from mESC, using an approach similar to ours. When these cells were co-cultured with PGCLC, reconstituted oocytes were formed that could undergo meiosis and produced live and healthy pups (Yoshino et al., 2021). The approaches we have developed could be exploited to examine the capability of these cells to support *in-vitro* spermatogenesis and oogenesis leading to the understanding of a range of fertility-related issues (Hayashi et al., 2011).

Our model is a powerful approach to understand testis-determination and differentiation in mouse and human as well as cellular and molecular effects of genes/variants associated with DSDs.

## Acknowledgements

This work is funded in part by a research grant from the European Society of Pediatric Endocrinology (to AB), and by the Agence Nationale de la Recherche (ANR; ANR-10-LABX-73 REVIVE to KM, ANR-17-CE14-0038-01 to KM, ANR-19-CE14-0022 to AB and ANR-19-CE14-0012 to AB). NG, AndB, RM and RLB were funded by the Francis Crick Institute. The Francis Crick Institute receives its core funding from Cancer Research UK (FC001107), the UK Medical Research Council (FC001107), the Wellcome (FC001107), and by the UK Medical Research Council (U117512772). We are grateful to the Flow Cytometry, Bioinformatics and Biostatistics as well as Advanced Sequencing Facilities of the Francis Crick Institute. The hiPSCs were generated by Phenocell, France. We thank Delphine Bohl and Stephanie Bigou at Institut du Cerveau - Hôpital Pitié Salpêtrière, Paris for their help with generation of gene edited hiPSC lines.

## Author contribution

CE, NG, KM, RLB, and AB conceived the experiments; CE, NG, AB, AC and AndB performed the experiments; RM analysed the mouse RNA-Seq data and performed the computational analyses. SG designed the microfluidic device and SG, CE, EF and AB optimized its protocols. PHC conducted the flow-cytometric analysis of hiPSCs. IM provided primary patient care. CE, NG, RLB, KM, AB collated the experimental data. The manuscript was written by AB, KM, RLB, CE and NG and reviewed by AndB and RM. All the authors read and agreed with the data being presented in the manuscript.

## Ethics declaration

The authors declare no competing interests.

