## Supplementary data- methods, figures and tables for "*In-vitro* cellular reprogramming to model gonad development and its disorders"

### Supplementary files

#### Materials and Methods

##### MICE

All animals were maintained with appropriate care according to the United Kingdom Animal Scientific Procedures Act 1986 and the ethics guidelines of the Francis Crick Institute.

##### Derivation of mouse ESC from blastocysts

XY Mouse ESC (TESCO-CFP; R26-rtTA, clone 7) were derived from the inner cell mass of blastocyst-staged embryos. TESCO-CFP heterozygous mice were bred with R26-M2rtTA homozygous mice (generated from Jax mouse line 016836- R26-M2rtTA; TetOP-H2B-GFP). E4.5 Blastocyst embryos were flushed in M2 media (M7167, Sigma) from the uterus and each embryo was placed on a 0.1% gelatin coated 24-well dish containing NDiff+2i/LIF media (described below). 7-14 days following seeding, a colony was formed and expanded. Cells were genotyped for sex chromosomes (McFarlane et al., 2013), TESCO-CFP (Gonen et al., 2018) and the R26-M2rtTA allele (Jax protocol https://www.jax.org/strain/016836). Primers used for genotyping are depicted in Table S1. Most of the differentiation work was carried using clone 7 which was XY, TESCO-CFP; R26-M2rtTA.

##### Mouse ESC culture

TESCO-CFP; R26-M2rtTA; XY ESCs were routinely cultured on 0.1% Gelatin (G9391, Sigma) coated dish with 2i/LIF media [NDiff medium (Y40002, Takara) supplemented with Penicillin-Streptomycin (P/S-15140-122, Invitrogen), L-Glutamine (L-Glu-25030-024, Invitrogen), Beta-mercaptoethanol (M6250, Sigma), 1 µM of PD0325901 (1408, Axon), 3 µM of CHIR99021 (1386, Axon), and 1000 U/ml of LIF (ESG1107, Millipore)].

##### Intermediate mesoderm differentiation of mouse ESCs

For differentiation, ESC cultured on 2i/LIF were seeded for 1 passage on dishes coated with Gel-MEF: 0.1% gelatin coating for 15 minutes followed by coating with MEF media for at least 2 hours [DMEM, 10% FCS, P/S, L-Glu]. ESCs were seeded into KSR media [Ad-DMEM/F12 (12634, Life technologies), 20% knock out serum replacement (10828010, Life technologies), P/S, L-Glu, Beta-mercaptoethanol and 1000 U/ml of LIF].

**For epiblast-like cells (EpiLCs) differentiation**, $2.5 \times 10^5$ cells were seeded on Gel-MEF coated 6-well dish wells. EpiLC differentiation was performed as previously described

(Hayashi et al., 2011). Briefly, EpiLCs were induced by culturing ESC in N2B27 minus Vitamin A (12587-010, Life technologies) medium containing 20 ng/ml activin A (338-AC, R&D), 12 ng/ml bFGF (233-FB, R&D) and 1% knockout serum replacement (KSR, 10828010, Life technologies) for 2.5 days (~60 hours).

Early mesoderm differentiation was induced by growing the cells as previously described (Bernardo et al., 2011; Brons et al., 2007) in a chemically defined medium (CDM) containing PVA (P8136, Sigma) and supplemented with 20 ng/ml bFGF, 10 μM LY294002 (1130, TOCRIS) and 10 ng/ml BMP4 (314-BP, R&D), named FLyB for 36 hours.

Intermediate mesoderm differentiation was performed for 3.5 days in CDM-PVA media supplemented with the following growth factors as described in the result section (IM1-5): FGF (5ng/ml), BMP4 (20ng/ml), RA (100nM, R2625, Sigma), CHIR99021 (0.3 μM), activin A (10ng/ml). IM3 which showed the best results contained FGF (5ng/ml), BMP4 (20ng/ml) and RA (100nM).

**IM1**: FGF (5ng/ml) + BMP (20ng/ml)

**IM2**: FGF (5ng/ml) + BMP (20ng/ml) + RA (100nM) + A (10ng/ml)

**IM3**: FGF (5ng/ml) + BMP (20ng/ml) + RA (100nM)

**IM4**: FGF (5ng/ml) + BMP (20ng/ml) + RA (100nM) + CHIR (0.3 μM)

**IM5**: FGF (5ng/ml) + BMP (20ng/ml) + RA (100nM) + A (10ng/ml) + CHIR (0.3 μM)

Following viral induction (described below), IM cells were cultured in a **"Sertoli media"** containing Ad-DMEM/F12 supplemented with P/S, L-Glu, HEPES, X1 B27 (GIBCO 17504-044) and nAC (Sigma A9165, 1.25 mM). The following growth factors were added: mEGF (Invitrogen PMG8043, 50 ng/ml), rhFSH (R&D 5925-FS, 22ng/ml), rmFGF9 (R&D 7399-F9, 50 ng/ml), Prostaglandin D2 (Cayman 12010, 500 ng/ml), testosterone (Sigma T1500, 1μM) and Activin A (R&D 338-AC, 50 ng/ml).

#### Lentivirus production

Lentiviruses encoding Nr5a1 and Dmrt1 were generated by transfecting HEK293T cells with the plasmids FUW-TetO-Nr5a1/Sf1 (Addgene 41081, gift from Y. Buganim) or FUW-TetO-Dmrt1 (Addgene 41083, gift from Y. Buganim) along with the packaging vectors pMD2.G (Addgene 12259) and psPAX2 (Addgene 12260) using XtremeHP transfection reagent (6366236001, Roche). A day after transfection media was replaced to minimal volume of IM3 media and viral particles were collected 48 hours post transfection. Supernatant containing

viral particles was filtered through 0.45 µm filter. Viral particles were supplemented with 2 µg/ml of polybrene (Sigma, H9268) and were used for infection of IM cells.
Following 3.5 days of IM differentiation, each 6 well was split into 2 new wells and viruses produced in IM media were added onto the newly seeded cells. Viruses were left on the cells for 24 hr after which media was changed to "Sertoli media" supplemented with 2 µg/ml Doxycycline (D9891, Sigma).

**Flow cytometry of mouse differentiated CFP positive cells**

CFP expressing early gonadal progenitors were FACS sorted 4 days following Dox addition. Cells were prepared into single cell suspension using trypsinization and filtered via a 30 µm filter (CellTrics, 04-004-2326). Sorting was performed on the BD *FACSARIA*™ III cell sorter using a 450 laser. CFP negative IM3 cell sample was used to set up the gating.

**RNA isolation, cDNA preparation and Quantitative Real-Time Polymerase chain reaction (qRT-PCR) for mouse differentiated cells**

Total RNA was extracted using Tri Reagent (Sigma, T9424) according to the manufacturer's protocol. RNA yield was quantified using a NanoDrop spectrophotometer (NanoDrop Technologies), and 1500 ng RNA was treated with RQ1 DNase (Promega) and used to synthesize cDNA using the SuperScript™ III Reverse Transcriptase kit (Invitrogen). qRT-PCR reactions were performed in duplicate using SYBR Green PCR master mix (Invitrogen) and 150 nM each of forward and reverse primers and analysed on the Applied Biosystems 7500 Real-Time PCR System (Thermo Fischer Scientific). Primers are listed in Table S2. Relative mRNA levels were determined by calculating $2^{-\Delta\Delta Ct}$ values relative to the normalizer gene PBGD. Relative gene expression is presented as the mean $2^{-\Delta\Delta Ct}$ values (error bars are SEM of the $2^{-\Delta\Delta Ct}$) for triple biological repeats.

**RNA-Sequencing**

Total RNA was extracted Tri Reagent as described above. Triplicate of RNA samples of ESC, EpiSC-like cells, M-like cells, IM-like cells (IM1-5) and CFP sorted Sertoli-like cells were used for RNA-Seq. RNA quality and quantity was analysed on the Qubit (Thermo Fischer Scientific) and Tapestation (Agilent). Libraries were prepared using the TrueSeq Library Prep kit V2 (Illumina) according to the manufacturer's instructions.
Sequencing was performed on the Illumina HiSeq 4000 system (Paired end, 75 bp).

#### Bioinformatic analysis

The mouse ESC differentiation RNA-Seq data has been deposited to the GEO under the accession number: GSE165133.

#### Data collection

Published RNA-Seq data from mouse embryonic gonad tissue relating to stages E10.5, E11.5, E12.5 and E13.5 were downloaded from the NCBI's Short Read Archive under accession: SRP076584 (Zhao et al., 2018). Published RNA-Seq data from sorted TESCO-CFP E15.5 Sertoli cells was downloaded from the NCBI's Short Read Archive under accession: SRP033562 (Maatouk et al., 2017).

#### Alignments and abundance estimation

Cutadapt v1.9.1 was used to trim adapter sequences from reads with the following options: -a AGATCGGAAGAGC –A AGATCGGAAGAGC -e 0.1 –q 10. Gene-level abundance estimates were calculated with RSEM v1.3.0 (Dobin et al., 2013), using STAR v2.5.2a (Li and Dewey, 2011) to align reads against the GRCm38 genome assembly with Ensembl release 86 transcript annotations. A copy of the code used for this step is freely available as a Nextflow pipeline here.

#### Data exploration and differential expression

Gene-level RSEM estimated counts and average transcripts lengths were imported into R 3.6.0 using the tximport function from the tximport package (Soneson et al., 2015). These were used to create a DESeqDataSet object for further analysis using the Bioconductor package DESeq2 (Anders and Huber, 2010). Data were normalised for differing library size using DESeq2's default method, leveraging the average transcript length information. Differential gene expression analysis between replicate groups was assessed using the default Wald test. Genes were called significant it they passed a combined filter of i) FDR<=0.01, ii) fold change >= +/-2, iii) base-mean >= 100 from the Wald test results and iv) a mean normalised read count of >= 100 in at least one of the tested replicate groups.

Principal Components Analysis (PCA) was used to assess the relationship between gene expression across samples using the PCAtools package's "pca" function (center=TRUE, scale=FALSE, removeVar=0.9). Data were first variance stabilised using DESeq2's "vst" function, which is roughly similar to putting the data on the log2 scale, while also dealing with the sampling variability of low counts. Only the top 10% most variant genes across selected samples were used to generate the visualisations.

While majority of the samples were generated *in-vitro* there are a number that were generated *in-vivo*, specifically: E10.5, E11.5, E12.5, E13.5 gonads and E15.5 sorted Sertoli cells. This difference in protocol appeared to dominate the first principal component (PC1, 47.47% variation) when combining both *in-vivo* and *in-vitro* data. A bi-plot of PC2 and PC3 better reflected the expected biology.

The *in-vitro/in-vivo* protocol specific effect was modelled and removed from the variance stabilised data using the limma package's "removeBatchEffect" function for the purposes of visualisation of combined *in-vitro* and *in-vivo* samples only. PCA analysis of the corrected data showed that PC1 was analogous to PC2 of the uncorrected data.

Sample similarity was assessed using a Poisson dissimilarity matrix constructed from the un-corrected normalised counts of all samples using the "PoissonDistance" function from the PoiClaClu package. All genes with a count greater than 0 in at least a single sample were included in the generation of the matrix unless otherwise stated.

Heatmaps were generated using the variance stabilised data. Data were additionally scaled per gene using a z-score to aid visualisation. Columns (samples) and rows (genes) were each hierarchically clustered using a "complete" clustering method on a set of Euclidean distances.

**Immunofluorescence staining for mouse differentiated cells**

Immuno staining was performed onto glass chamber slides. Cells were fixed for 10 min in 4% PFA (Sigma, P6148**)** at RT. Blocking was performed in 5% Donkey serum (Sigma, D9663) in PBS + 0.1% Triton (PBST) solution for 1 hr at RT. Primary antibodies were incubated in PBST supplemented with 1% Donkey serum O/N at 4ºC. Staining was performed using the following primary antibodies: Rabbit polyclonal anti-WT1 antibody (1:100, Santa Cruz, sc192), Goat polyclonal anti-GATA4 antibody (1:200, Santa Cruz, sc1237), Goat anti-BRACHYURY (1:150, R&D systems, AF8025). 3 washes in PBST were performed and secondary antibodies were incubated in PBST supplemented with 1% Donkey serum for 1 hr at RT. Secondary antibodies used were Donkey anti-rabbit/goat Alexa Fluor 568 (1:500, Invitrogen) and Donkey anti-goat Alexa Fluor 488 (1:500, Invitrogen). All immunofluorescence slides were also stained with 4',6-diamidino-2-phenylindole (DAPI, Molecular Probes), to visualize nuclear DNA. Images were taken on the Zeiss LSM710 confocal microscope.

#### HUMAN

##### Human induced Pluripotent Stem Cell (hiPSC) lines

Peripheral blood was collected from two individuals. First, a 46,XY girl carrying a *de novo* heterozygous p.Arg313Cys pathogenic variant in *NR5A1*, identified by exome sequencing that causes a complete lack of testis determination (46,XY complete gonadal dysgenesis, (Mazen et al., 2016)). The second sample was from her brother, a healthy 46,XY male, who did not carry the *NR5A1* variant. Peripheral blood mononuclear cells (PBMC) from these two samples were reprogrammed into induced pluripotent stem cells (Phenocell, Grasse). Briefly, PBMCs were cultured in complete StemSpan™ SFEM II medium (#09605; StemCell Technologies) were reprogrammed with the Epi5™ Episomal iPSC Reprogramming Kit (#A15960, ThermoFisher Scientific) according to manufacturer's instructions. The reprogramming vectors were introduced by nucleofection and the transfected cells were harvested and plated on Laminin 521-coated P6 culture dish in ReproTeSR™ medium (#05926; StemCell Technologies). Medium was changed daily until iPSC colonies appeared (10-12 days), then diluted 1/1 with mTEeSR™1 (#85850; StemCell Technologies) until colonies were large enough to pick and expand. Colonies are selected on morphological criteria, isolated, and further amplified in mTEeSR™1 over 10-12 passages to perform quality control by Karyotyping, Array CGH and exome sequencing. A third iPSC line was bought from Phenocell. Briefly, this line was reprogrammed from PBMC obtained from an 46,XX female donor. A fourth line of iPSC was generated after correction of the *NR5A1* mutation, using CRISPR/CAS9, in the iPSCs derived from 46,XY girl (CELIS platform, Institut du Cerveau et de la Moëlle, ICM, la Pitié-Salpétriere, PARIS).

##### Culture of hiPSC

hiPSCs were cultivated in feeder-free mTeSR$^{TM}$Plus medium (#100-0276, StemCell Technologies) on qualified "human Embryonic StemCell" (hESC) matrigel (#354277, Corning). Thiazovivin was added for 48 hours to the medium to prevent spontaneous differentiation (#130-106-542, Miltenyi Biotec) at thawing and passaging. When the colonies reached 70-80% of confluence, cells were split into clumps or aggregates with the ReLeSR (#05872, StemCell Technologies) medium and, depending on the confluence, clumps were diluted at 1:10 or 1:20 into a new freshly Matrigel coated flask.

**Differentiation of hiPSC**

24 hours before differentiation, colonies of iPSC (80% of confluence) were dissociated into single cells solution, using the Gentle Cell Dissociation reagent (#07174, StemCell Technologies), and seeded on Matrigel coated flasks and plates (T25cm$^2$ flask, 6-well plate and chamber slide) with a high density of cells (1:2 dilution). Similar to murine pluripotent cells, hiPSCs were subjected to serial differentiation in conditioned medium with minor modifications of the medium composition. The basal medium used for subsequent steps was Chemically Defined Medium (CDM-PVA) containing 250ml of advanced DMEM (#31331028, ThermoFisher Scientific), 250ml of Iscove's Modified Dulbecco's Media (IMDM, #31980030, ThermoFisher Scientific), 0.1% of cold water soluble polyvynilic alcohol (#P8136, MerckMillipore), 5ml of penicillin-streptomycin (#15140122, 10 000U/ml, ThermoFisher Scientific), 5ml of concentrated lipids (11905031, 1:100, ThermoFisher Scientific), monothioglycerol (MTG, 20ul, water miscible 0.1M, #M6145, MerckMillipore) and 300ul of transferrin (#1065220200, water soluble, MerckMillipore). For mesodermal induction, cells were incubated for 36 hours in FlyB medium (CDM-PVA with bFGF (20ng/ml; #233-FB, R&D), Ly294002 (10µM; Pi3K inhibitor, #L9908, MerckMillipore) and BMP (10ng/ml; #214-BP, R&D)). This was followed by directed differentiation towards intermediate mesoderm using IM3 medium, composed of CDM-PVA with bFGF (5ng/ml), BMP (20ng/ml) and retinoic acid (RA, 100nM; #R2625, MerckMillipore) for 48 hours, with a change of medium at 24 hours. To induce differentiation toward supporting cell lineages, the medium was changed to supporting Medium that is composed of 500ml of Advanced DMEM (#12634010, ThermoFisher Scientific), 5ml of Penicillin/ Streptomycin, 5ml of Insulin, Transferrin, Selenium (ITS, 100X #12097549, ThermoFisher Scientific) and 50ul of EGF (20 ng/mL; human recombinant #ab9697, Abcam). The differentiating cells were cultured in supporting medium, with the change of medium every 2-3 days until spontaneous tubular structures appear.

**Flow cytometry of differentiated hiPSC**

Once defined structures are visible the cells are dissociated into single cells. For cell sorting, 1x10$^7$ cells were incubated with rabbit anti-CLAUDIN11 antibody (dilution 1:100, #36-4500, ThermoFisher Scientific) followed by anti-rabbit IgG associated with either Alexa 488 (#A11034, ThermoFisher Scientific) or 594 (#A11037, ThermoFisher Scientific). The stained cells were collected in FACS tube with cell strainer (#352235, Corning) and examined using MoFLO Astrios “Beckman Coulter” with Summit v62 (Beckman Coulter). After negative selection for Alexa 488 or 594 remaining gated cells were collected in Eppendorf tube containing 500µl of supporting medium. On an average CLAUDIN11 positive cells represented between 4-12% of the initial population depending on the differentiation efficiency. Sorted cells were seeded in a well of a 12 well-plate coated with non-qualified Matrigel (#354230, Corning), and cultured for several weeks to let the cells recover from the stress of sorting and grow.

**Immunofluorescence staining for undifferentiated hiPSCs and differentiated cells**

Undifferentiated hiPSC and differentiated cells were cultured on chamber-slides, and stained for OCT4/POU5F1, SOX9, FOXL2, CLAUDIN11 and DMRT1 proteins according to the protocol described elsewhere (Eozenou et al., 2020). Briefly, after fixation in 4% PFA, permeabilization and blocking of non-specific epitopes, cells were incubated with the primary antibody (diluted in 3% BSA in PBS) in a humid chamber overnight at +4°C. The following dilutions of primary antibodies were used: anti-SOX9 (1:100, #14-9765-82, ThermoFisher Scientific), anti-OCT4 (1:100, #ab19857, Abcam), anti-FOXL2 (1:100, #ab5096, Abcam), anti-CLAUDIN11 (1:100, #36-4500, Life Technologies) and anti-DMRT1 (1:100, #ab126741, Abcam). After 16 h of incubation, cells were washed three times with 1xPBS for 5 min each. This was followed by incubation with the secondary antibody in 3% BSA for 1h at room temperature in the dark. Depending on the primary antibody, the following secondary antibodies were used- Goat anti-Rabbit IgG (H+L) Secondary Antibody, Alexa Fluor® 594 conjugate (1:1000, #A11037, Life Technologies), Goat anti-mouse IgG (H+L) Secondary Antibody, Alexa Fluor® 488 conjugate, (1:1000, #A11029 Life Technologies), Donkey anti-goat IgG (H+L) secondary antibody, Alexa Fluor® 488 conjugate, (1:1000, #ab150129, Abcam). Cells were washed three times with 1xPBS for 5 min each in the dark and incubated with DAPI diluted in PBS for 15min in the dark (1:2000, #62248, ThermoFisher Scientific). After three washes in PBS, slides were mounted using ProLong® Gold Antifade Mountant with DAPI (#P36931, ThermoFisher Scientific). Images were obtained with a Leica Microsystems DMI4000B microscope at 40x, 63x and 100X (with oil) magnifications.

**Quantitative Real-Time Polymerase chain reaction (qRT-PCR) for undifferentiated hiPSCs and derivatives**

Total RNA was extracted from undifferentiated hiPSC and cells during the course of differentiation, using TRIzol reagent (#15596026, ThermoFisher Scientific). RNA yield was quantified with a NanoDrop spectrophotometer (NanoDrop Technologies), and 1000 ng RNA was used to synthesize complementary DNA (cDNA) with the Quantitect Reverse Transcription Kit (#205311, QIAGEN) as per manufacturer's recommendations. cDNA was diluted 1/25 prior to the qPCR. qPCR was performed using TaqMan Universal Master Mix II, with UNG (#4440038, Applied Biosystems) on a StepONEPlus qPCR machine (Applied Biosystems). The Following TaqMan probes (Applied Biosystems) were used; RPL19: #Hs02338565_gH; SOX9: #Hs01001343_g1; DMRT1: #Hs00232766_m1; NR5A1: #Hs00610436_m1; WT1: #Hs01103751_m1; FOXL2: #Hs00846401_s1; FGF9: #Hs00181829_m1; OCT4/POU5F1: #Hs04260367_gH; NANOG: #Hs02387400_g1; BRACHYURY/T: #Hs00610080_m1; NR2F2: #Hs00819630_m1; OSR1: #Hs01586544_m1; GATA4: #Hs0171403_m1; RSPO1: #Hs00543475_m. Relative mRNA levels were determined by calculating $2^{-\Delta\Delta Ct}$ values relative to the 18S rRNA normalizer gene (RPL19). Relative gene expression is presented as the mean $2^{-\Delta\Delta Ct}$ values (error bars are SEM of the $2^{-\Delta\Delta Ct}$).

**Statistics**

Statistical analyses were carried out using GraphPad Prism 8 software (GraphPad). Quantitative data were subjected to a one-way ANOVA (or student t-test) followed by Bonferroni comparison.

**Measurement of AMH concentrations**

AMH (Anti-Müllerian Hormone) was measured by a one-step sandwich enzyme-linked immunosorbent assay (Access AMH, Beckman Coulter Company, Marseille, France (Pearson et al., 2016)). AMH is sandwiched between two anti-AMH monoclonal antibodies, one conjugated to alkaline phosphatase, the other coated with paramagnetic particles. A Lumi-Phos 530 chemiluminescent developer was used to read the light output proportional to the concentration of AMH in the sample relative to recombinant human AMH that was used as standard. The intra-assay coefficients of variation range from 1.41% to 3.3% and the inter-assay coefficients of variation from 3.04% to 5.76%.

#### Soft Matrigel substrates

Organoid grade Matrigel (#354263, Corning) was diluted to 50% (v/v) in supporting medium. 50-µl domes of Matrigel were cast onto a well of chilled 12-well plate and left for gelation for 30 min at 37°C. 50µl of concentrated cells ($1x10^6$ cells/ml) were pippeted onto the domes slowly and incubated at 37°C for 30min, after which 50µl of supporting medium was added carefully, and the plates were incubated for 48-72 hours. Cells self-aggregate on the top of the dome and make 3D structure and excess cells grow as a layer at the bottom of the dome. Images were obtained with a Leica Microsystems DMI4000B microscope at 40x, 63x and 100X (with oil) magnifications.

#### GONAChip microfluidic device and migration assay

GONAChip was composed of three channels (one central for the Matrigel flanked by two media channels) with gaps in the walls separating the channel in order to ensure a proper medium gel/medium interface. The proper confinement of the Matrigel in the central channel was ensured by hydrophobic pinning. Chips were produced by means of photo- and soft-lithography as described in Jeon *et al.* (Jeon et al., 2015). Briefly a master mould of 160 um in hight was created by means of photolithography. Replication of the device were performed with poly-dimethyl-siloxane (PDMS, Silgard 184; Dow Chemical). Finally, we used oxygen plasma to bond the PDMS slabs onto #1.5 Glass coverslips. Prior to use we performed a 20 min sterilization step in a UVO cleaner (Jelight, CA, US). Organ grade Matrigel (#354263, Corning) was diluted with advanced DMEM medium to have 50% (v/v) Matrigel (50µl of medium into 50µl of Matrigel). The central channel was filled with 1.3µl of this solution. After gelation at 37°C in the incubator, media channels were filled with supporting medium. Confluent sorted cells were detached, counted, and concentrated at $1x10^6$ cells/ml. Few µl of cell suspension was introduced in each chip. Chips were flipped to 90° to allow cells to sediment on the Matrigel channel side. After washing the chips were left in standard cell culture incubator (37°C, 5% CO2). When performing Time-lapse imaging, seeded GONAChips were placed under an inverted microscope equipped with a temperature, humidity, and CO2 control system. (Inverted Z1 Axio Observer, ZEISS). Pictures were taken every 15 min for 70 hours. Images are processed using the ZENlite Software (ZEISS) to extract videos and images.

**Correction of *NR5A1* p.Arg313Cys by CRISPR/CAS9**

$1x10^6$ hiPSCs were nucleofected with RNP complex (225 pmol of each RNA crRNA; #Alt-R® CRISPR-CAS9 crRNA, IDT; GCTGGACCTGGC**a**GTAGATG (The target site in the sequence is specified by the small bold letter), tracrRNA-ATTO+; #Alt-R® CRISPR-CAS9 tracrRNA, IDT) and 120 pmol of Cas9 protein; #Alt-R® S.P. Hifi CAS9 nucelase 3 NLS, IDT; #Alt-R® CAS9 electroporation enhancer, IDT) and HDR template (500 pmol ssODN; TGCCCGGTGACCAGCAGGATGCTGCCCTCCTTGCCGTGCTGGACCTGGC**g**GTAGA TGTGATCGAACACCAGCAGCTCGCTCCAGCAGTTCTGCAGCAGCG (The inserted change in the sequence is specified by the small bold letter); Ultramer DNA oligo (ssODN repair template), IDT). 24hrs later, ATTO+ transfected iPSCs were sorted by FACS and plated at very low density (10 cell/$cm^2$) on Ln521 (#77003, StemCell Technologies) with CloneR supplement (#05888, StemCell Technologies) for clonal selection. One week later, hiPSC clones were picked under a stereomicroscope and cultured on Ln521 in 96 well plates. When confluent, iPSC clones were duplicated for cryopreservation and DNA extraction. Clones were then analysed by PCR (iCS-digital™ PSC – 24 probes) to assess the genomic integrity of the stem cell lines both before and after correction by CRISPR/CAS9 modification (Stem Genomics, Montpellier, France).

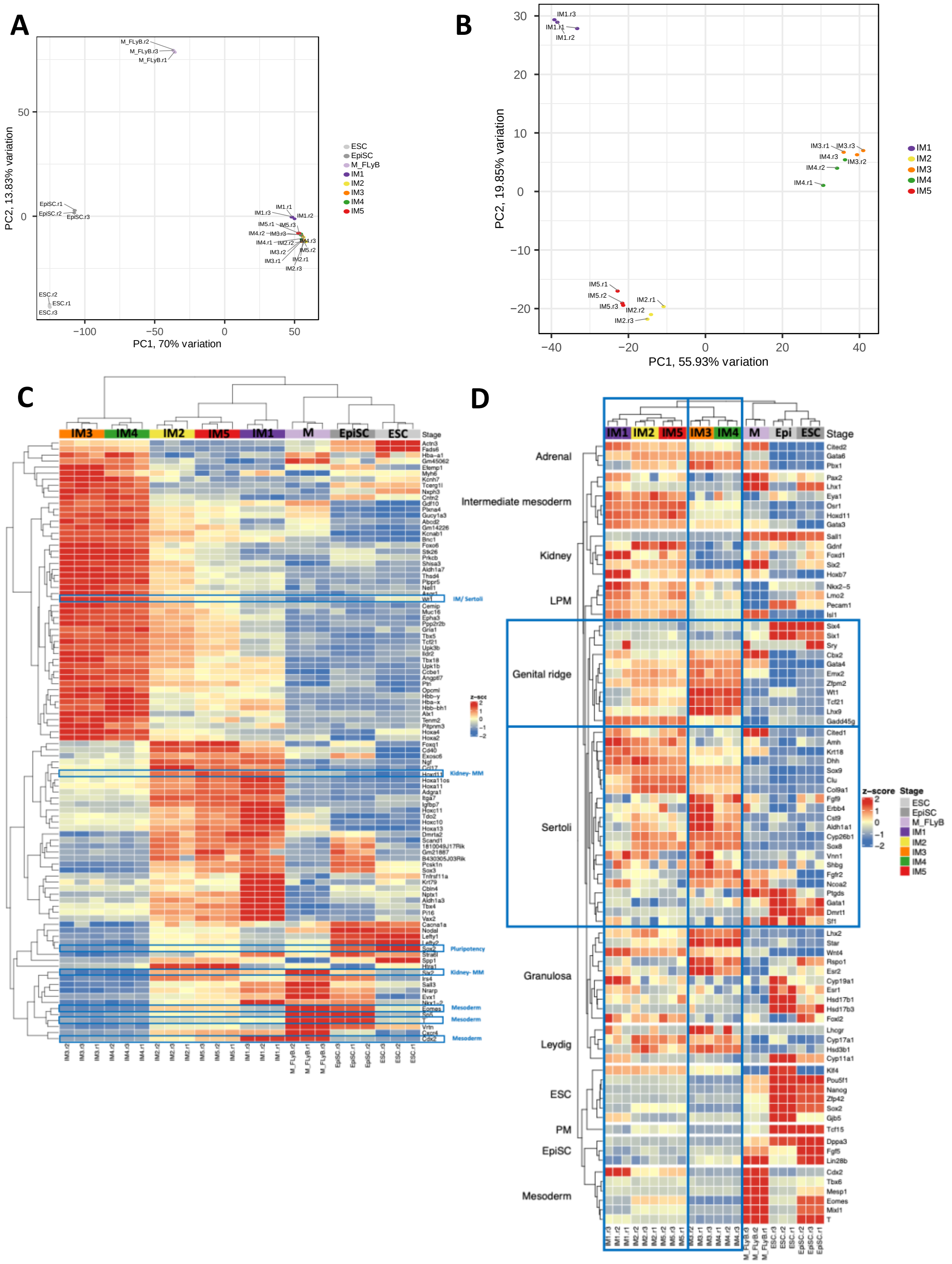


**Supplementary Figure 1. Transcriptomic analysis of the *in vitro*-differentiated cells** (A) Principal component analysis (PCA) of selected *in vitro* differentiated samples based on normalised mRNA expression level after batch correction. The top 2 PCs are shown. 70% of the variation is explained by PC1 and separates intermediate mesoderm samples from the other differentiation time points analysed. Three biological replicates were analysed for each sample type. (B) Principal component analysis (PCA) of the five different IM samples (IM 1-5) based on normalised mRNA expression level after batch correction. The top 2 PCs are shown. (C) Heatmap of the 100 most differentially expressed genes (50 most upregulated / 50 most down regulated) between IM1/2/5 and IM3/4. Markers of the various lineages are depicted by blue rectangles and the tissue they represent is denoted on the right. Three biological replicates were analysed for each sample type. (D) Heatmap of selected genes which represent known markers of the following lineages: Embryonic stem cells (ESC), Epiblast stem cells (EpiSC), Mesoderm (M), Intermediate mesoderm, Lateral plate mesoderm (LPM), Paraxial mesoderm (PM), Genital ridge, Sertoli cells, Leydig cells, Granulosa cells, Kidney and Adrenal. Three biological replicates were analysed for each sample type.

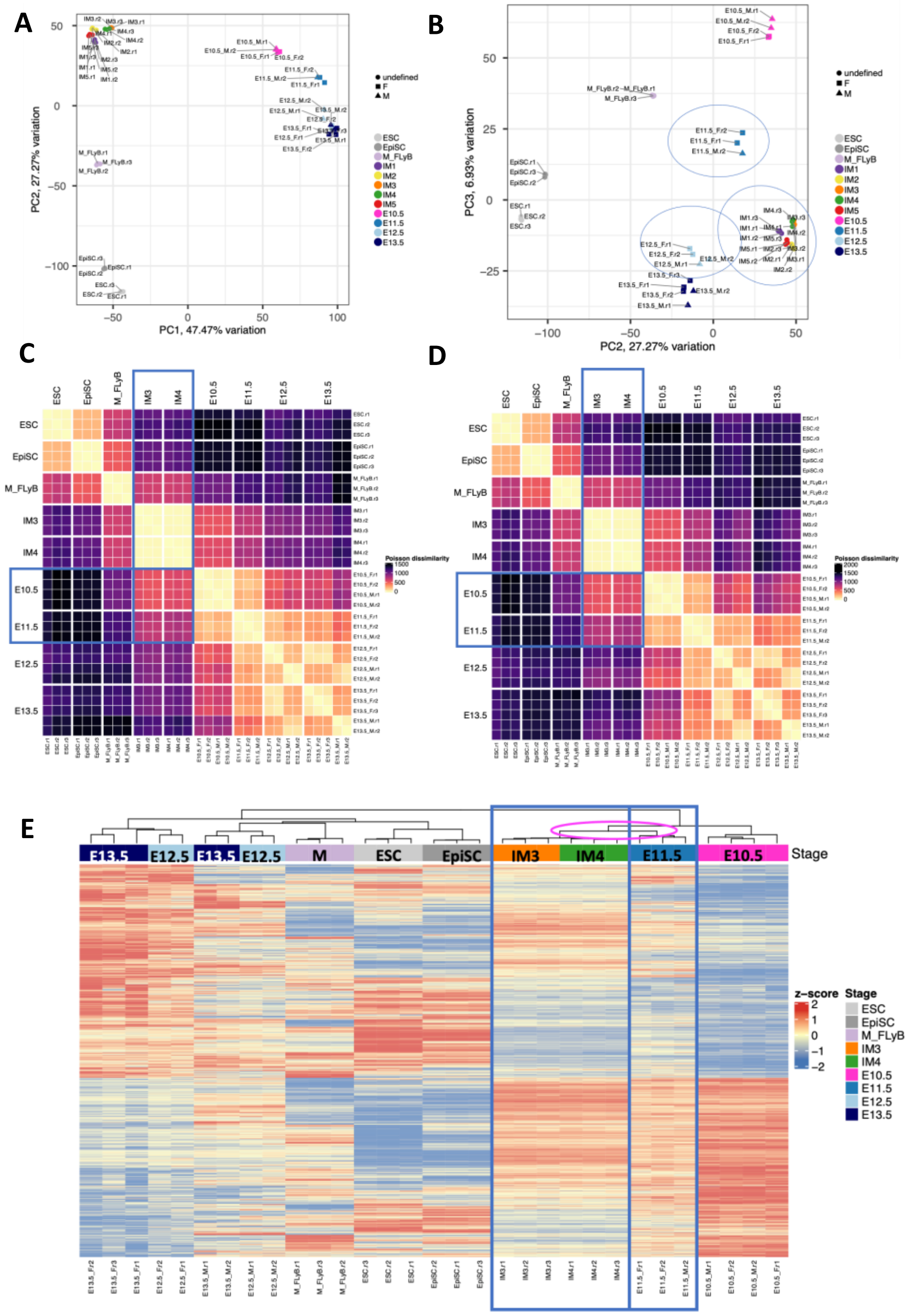


**Supplementary Figure 2. Transcriptomic comparison of *in vitro*- and *in vivo*-derived gonadal cells** (A) Principal component analysis of selected *in vitro* differentiated and *in vivo* gonad samples (Zhao et al., 2018) based on normalised mRNA expression level without batch correction. The top 2 PCs are shown. Three biological replicates were analysed for each sample type. Without data correction the PC1/PC2 separates between the *in vitro*-derived (left side) and *in vivo*-derived (right side) samples, while the PC2/PC3 (B) allows the temporal positional determination of IM *in vitro* derived cells within the *in vivo* samples analysed. (C) Heatmap of Poisson dissimilarity scores showing transcriptional similarities between samples. Dissimilarity scores were calculated from a normalised abundance matrix of genes differentially expressed between E10.5 XY and E13.5 XY gonads (Zhao et al., 2018). Blue boxes indicate the similarity between IM3/4 and E10.5-E11.5 gonadal cells. Dark purple denotes dissimilarity while bright yellow denotes high similarity. (D) Heatmap of Poisson dissimilarity scores showing transcriptional similarities between samples. Dissimilarity scores were calculated from a normalised abundance matrix of genes differentially expressed between E10.5 XX and E13.5 XX gonads (Zhao et al., 2018). Blue boxes indicate the similarity between IM3/4 and E10.5-E11.5 gonadal cells. (E) Heatmap of genes most differentially expressed between the E10.5 and E13.5 XX female gonads following batch correction (Zhao et al., 2018). IM3/IM4 cluster closely to the E11.5 *in vivo* gonadal cells. Gene-level expression across samples is shown as a z-score running from red (high) to blue (low). Three biological replicates were analysed for each sample type.

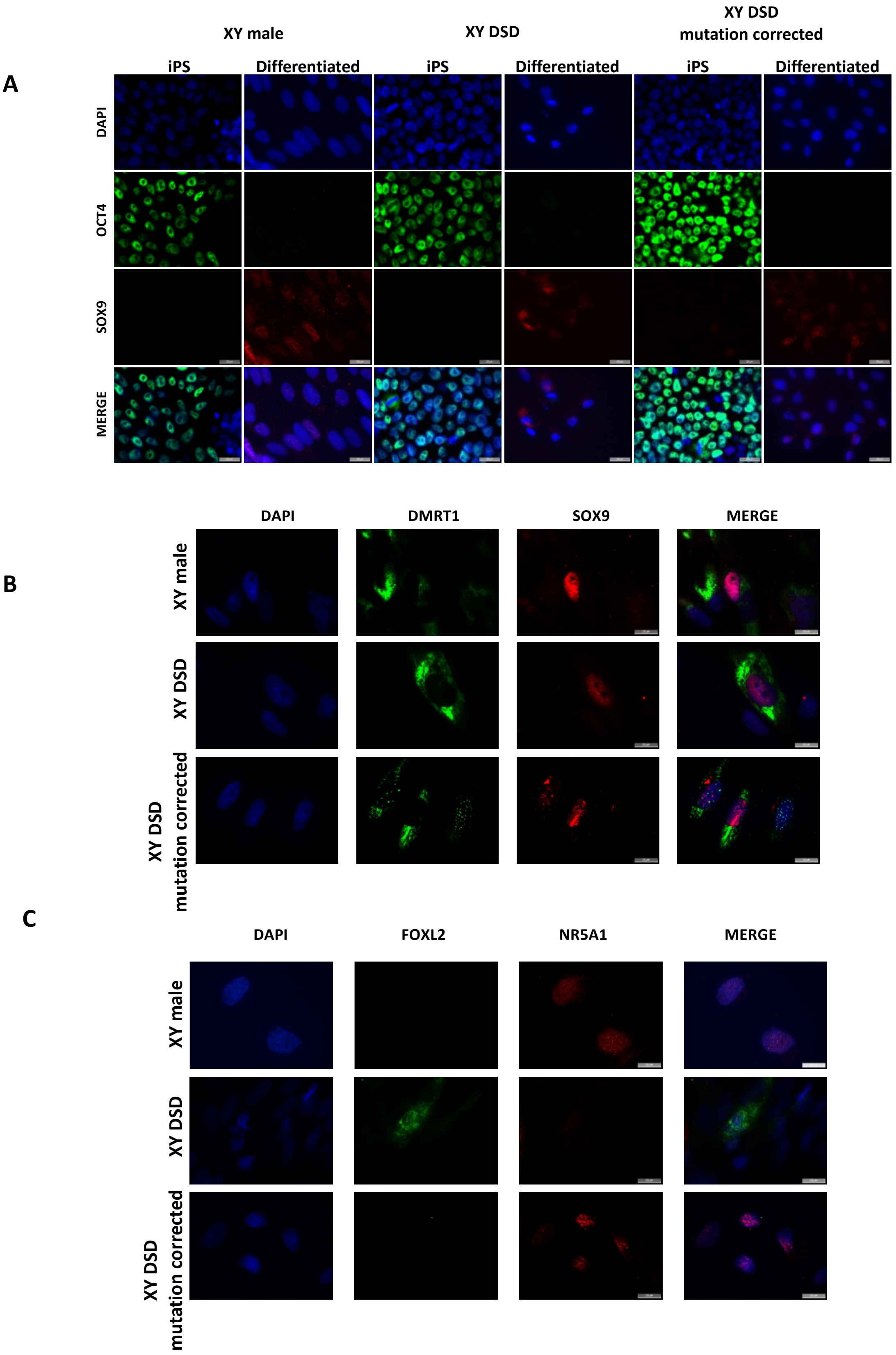


**Supplementary Figure 3**

Immunocytochemistry for OCT4 (A), SOX9 (A &B), DMRT1 (B), NR5A1 (C) and FOXL2 (C) expression in hiPSCs and somatic cells of the gonad derived from 46,XY healthy male,

46,XY DSD and NR5A1 CRISPR/CAS9 corrected 46,XY DSD cells following the differentiation protocol (bar = 20 μ)

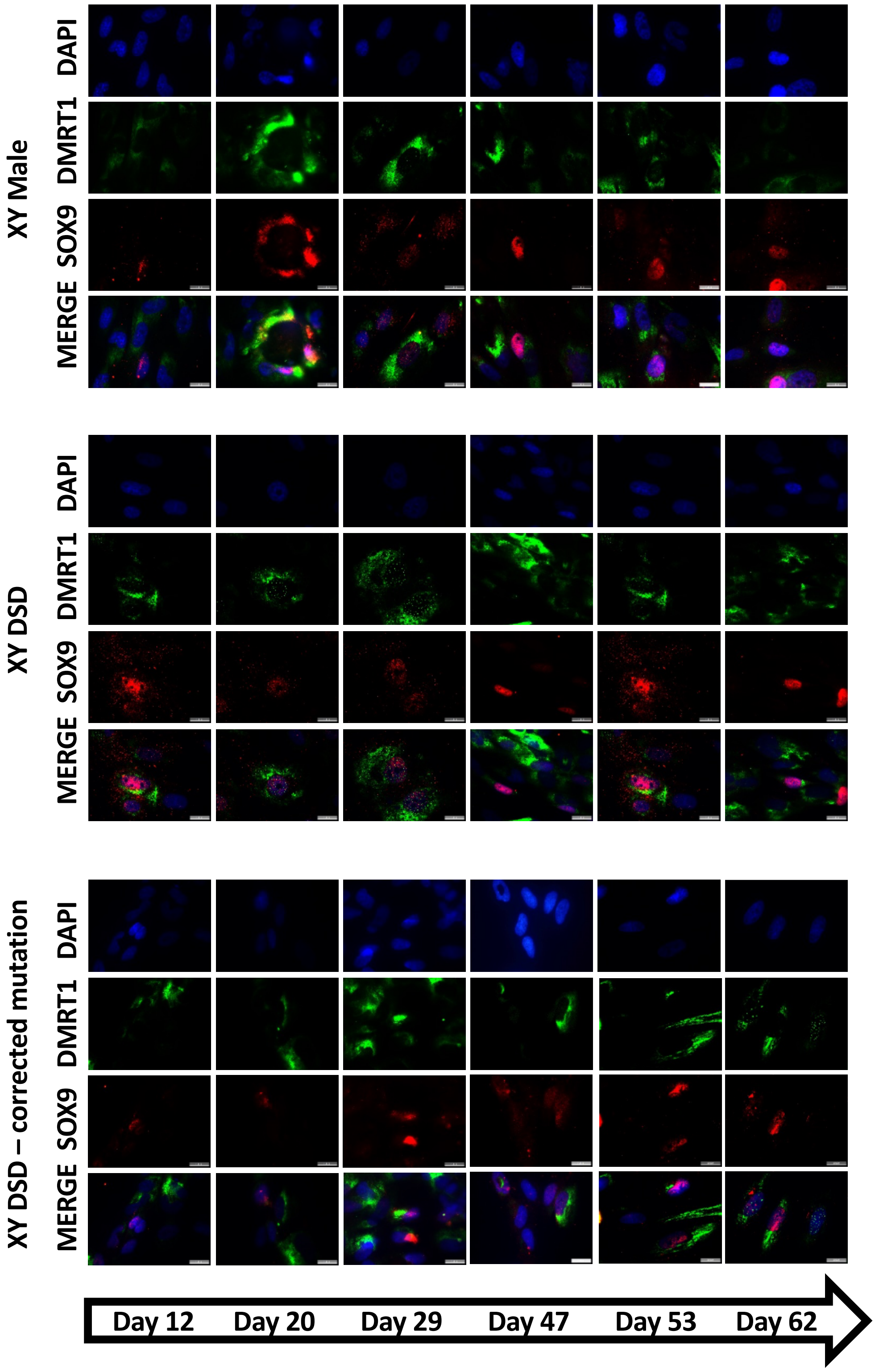


**Supplementary Figure 4**

Immunocytochemistry for SOX9 and DMRT1 expression in hiPSC-derived somatic cells of the gonad from 46,XY healthy male, 46,XY DSD and NR5A1 CRISPR/CAS9 corrected 46,XY DSD cells after sequential culture in defined media for 62 days (bar = 20 μ).

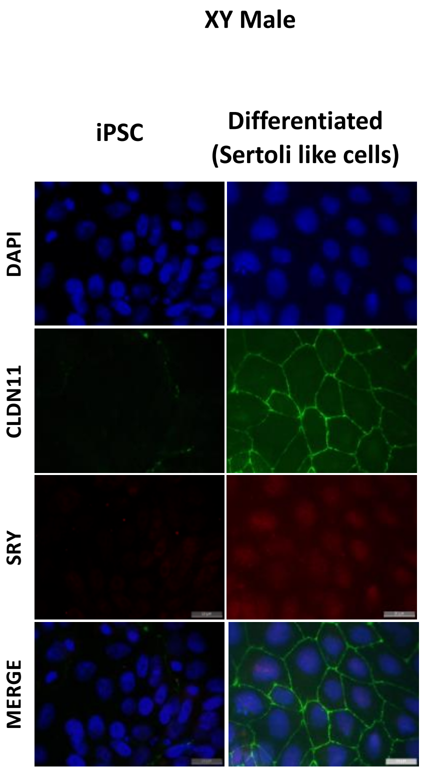


**Supplementary Figure 5**
Immunocytochemistry for SRY and CLAUDIN11expression in Sertoli-like cells derived from hiPSC-derived cells from a 46,XY healthy male (bar = 20 µ).

| Table S1. Primers for genotyping mice | | | |
|---|---|---|---|
| **Primer name** | **Description** | **Sequence 5' to 3'** | **Product and size** |
| CFP-F | CFP | CGTGACCACCCTGACCTGG | F+R: 400 bp product |
| CFP-R | CFP | GTGGCGGATCTTGAAGTTGG | |
| oIMR8052 | R26-M2rtTA_R_Mutant_SA site | GCGAAGAGTTTGTCCTCAACC | All 3 together: Mutant = 300 bp |
| oIMR8545 | R26-M2rtTA_F_R26_Common | GAAAGTCGCTCTGAGTTGTTAT | Heterozygote = 300 bp and 603 bp |
| oIMR8546 | R26-M2rtTA_R_R26 | GGAGCGGGAGAAATGGATATG | Wild type = 603 bp |
| Sex-F | X/Y chromosome | GATGATTTGAGTGGAAATGTGAGGTA | McFarlane et al., 2013 |
| Sex-R | X/Y chromosome | CTTATGTTTATAGGCATGCACCATGTA | |

| Table S2. Primers for Real-time quantitative RT-PCR | | | |
|---|---|---|---|
| **Primer name** | **Gene** | **Marker of** | **Sequence 5' to 3'** |
| mKLF4_F | *Klf4* | *ESC* | GACTAACCGTTGGCGTGA |
| mKLF4_R | | | GCCACTCTCCAGGTCTGTG |
| mFGF5_F | *Fgf5* | *EpiSC* | ACTCCATGCAAGTGCCAAAT |
| mFGF5_R | | | TCGGCCTGTCTTTTCAGTTC |
| mBra_F | *Bra (T)* | *Mesoderm* | AAGAACGGCAGGAGGATGT |
| mBra_R | | | TCACGAAGTCCAGCAAGAAA |
| mTbx6_F | *Tbx6* | *Mesoderm* | ACCGCTACCCTGATTTGGATA |
| mTbx6_R | | | AGATGGGAGAAGGGGCAAAG |
| mLhx1_F | *Lhx1* | *Intermediate mesoderm* | CCCATCCTGGACCGTTTCC |
| mLhx1_R | | | CGCTTGGAGAGATGCCCTG |
| mOsr1_F | *Osr1* | *Intermediate mesoderm* | GACCGCGGCGGAACAAGATA |
| mOsr1_R | | | CACTGTGGGCAGGCCATTCA |
| mPax2_F | *Pax2* | *Intermediate mesoderm* | AAGCCCGGAGTGATTGGTG |
| mPax2_R | | | CAGGCGAACATAGTCGGGTT |
| mWt1_F | *Wt1* | *Intermediate mesoderm/ Genital ridge* | TTGAATGCATGACCTGGAATCA |
| mWt1_R | | | TTCCCTTTAAGGTAGCTCCTAGGTT |
| mLhx9_F | *Lhx9* | *Genital ridge* | ACCAGCAGCCTTATCCACCTTCACAG |
| mLhx9_R | | | TGTAATGCCCCAAGATTTGTTCTCCC |
| mCbx2_F | *Cbx2/M33* | *Genital ridge* | GGCTGGTCCTCCAAACACAA |
| mCbx2_R | | | CCCTGGGTCTCTTGCCTCT |
| mPod1_F | *Pod1* | *Genital ridge* | CTCCCTGAAAGTGGACTCCAA |
| mPod1_R | | | CGGGCTTTTCTTAGTGGGC |
| mFog2_F | *Fog2* | *Genital ridge* | ACCAGGAGAGCTAGAAGTGTTT |
| mFog2_R | | | GGACCTGAGCCTTCGTCTT |
| mGata4_F | *Gata4* | *Genital ridge/ Sertoli cells* | CCCCAATCTCGATATGTTTGATG |
| mGata4_R | | | TTGACACACTCTCTGCCTTCTGA |
| mSox9_F | *Sox9* | *Sertoli cells* | AAGAAAGACCACCCCGATTACA |
| mSox9_R | | | CAGCGCCTTGAAGATAGCATT |
| mSf1_F | *Nr5a1/Sf1* | *Genital ridge/ Sertoli cells* | CCTCGATGTGAAATTCCTGAACA |
| mSf1_R | | | TCCTGGGCGTCCTTTACG |
| mDmrt1_F | *Dmrt1* | *Sertoli and Germ cells* | GGAGTCTCCCAGCACCTTACG |
| mDmrt1_R | | | TCTGCCACTGGTTTCCAGTCT |
| mPBGD_F | *PBGD* | *HK gene* | CCTGGCATACAGTTTGAAATCAT |
| mPBGD_R | | | TTTTTCCAGGGCGTTTTCT |

**Table S3. Gene list of selected markers of the various lineages used to compose Figure S1D**

| Tissue | Common name/ synonyms | gene_id | official gene_name | Comments |
|---|---|---|---|---|
| Adrenal | Cited2 | ENSMUSG00000039910 | Cited2 | |
| Adrenal | GATA6 | ENSMUSG00000005836 | Gata6 | |
| Adrenal | Pbx1 | ENSMUSG00000052534 | Pbx1 | |
| EpiSC | Lin28b | ENSMUSG00000063804 | Lin28b | |
| EpiSC | FGF5 | ENSMUSG00000029337 | Fgf5 | |
| EpiSC | Dppa3/ Stella | ENSMUSG00000046323 | Dppa3 | |
| ESC | Gjb5 | ENSMUSG00000042357 | Gjb5 | |
| ESC | KLF4 | ENSMUSG00000003032 | Klf4 | |
| ESC | Nanog | ENSMUSG00000012396 | Nanog | |
| ESC | Sox2 | ENSMUSG00000074637 | Sox2 | |
| ESC | Pou5f1/Oct4 | ENSMUSG00000024406 | Pou5f1 | |
| ESC | Zfp42/Rex1 | ENSMUSG00000051176 | Zfp42 | |
| Genital.ridge | Emx2 | ENSMUSG00000043969 | Emx2 | |
| Genital.ridge | GADD45g | ENSMUSG00000021453 | Gadd45g | |
| Genital.ridge | GATA4 | ENSMUSG00000021944 | Gata4 | |
| Genital.ridge | Lhx9 | ENSMUSG00000019230 | Lhx9 | |
| Genital.ridge | Six1 | ENSMUSG00000051367 | Six1 | |
| Genital.ridge | Six4 | ENSMUSG00000034460 | Six4 | |
| Genital.ridge | Sry | ENSMUSG00000069036 | Sry | |
| Genital.ridge | WT1 | ENSMUSG00000016458 | Wt1 | |
| Genital.ridge | Cbx2/M33 | ENSMUSG00000025577 | Cbx2 | |
| Genital.ridge | FOG2/ Zfpm2 | ENSMUSG00000022306 | Zfpm2 | |
| Genital.ridge | Pod1/Tcf21 | ENSMUSG00000045680 | Tcf21 | |
| Granulosa | Cyp19a1 | ENSMUSG00000032274 | Cyp19a1 | |
| Granulosa | Esr1 | ENSMUSG00000019768 | Esr1 | |
| Granulosa | Esr2 | ENSMUSG00000021055 | Esr2 | |
| Granulosa | Foxl2 | ENSMUSG00000050397 | Foxl2 | |
| Granulosa | Hsd17B1 | ENSMUSG00000019301 | Hsd17b1 | |
| Granulosa | Hsd17B3 | ENSMUSG00000033122 | Hsd17b3 | |
| Granulosa | Lhx2 | ENSMUSG00000000247 | Lhx2 | |
| Granulosa | Star | ENSMUSG00000031574 | Star | |
| Granulosa | Wnt4 | ENSMUSG00000036856 | Wnt4 | |
| Granulosa | Rspo1 | ENSMUSG00000028871 | Rspo1 | |
| Intermediate.Mesoderm | EYA1 | ENSMUSG00000025932 | Eya1 | Posterior IM |
| Intermediate.Mesoderm | GATA3 | ENSMUSG00000015619 | Gata3 | Anterior IM |
| Intermediate.Mesoderm | HOXD11 | ENSMUSG00000042499 | Hoxd11 | Posterior IM |
| Intermediate.Mesoderm | Lhx1 | ENSMUSG00000018698 | Lhx1 | Anterior IM |
| Intermediate.Mesoderm | Osr1 | ENSMUSG00000048387 | Osr1 | |
| Intermediate.Mesoderm | Pax2 | ENSMUSG00000004231 | Pax2 | |
| Intermediate.Mesoderm | WT1 | ENSMUSG00000016458 | Wt1 | |
| Kidney | FoxD1 | ENSMUSG00000078302 | Foxd1 | |
| Kidney | GATA3 | ENSMUSG00000015619 | Gata3 | |
| Kidney | GDNF | ENSMUSG00000022144 | Gdnf | Metanephric mesenchyme |
| Kidney | HoxB7 | ENSMUSG00000038721 | Hoxb7 | |
| Kidney | HoxD11 | ENSMUSG00000042499 | Hoxd11 | Metanephric mesenchyme |
| Kidney | PAX2 | ENSMUSG00000004231 | Pax2 | Ureteric epithelium |
| Kidney | SALL1 | ENSMUSG00000031665 | Sall1 | |
| Kidney | Six1 | ENSMUSG00000051367 | Six1 | Metanephric mesenchyme |
| Kidney | Six2 | ENSMUSG00000024134 | Six2 | Metanephric mesenchyme |
| Kidney | WT1 | ENSMUSG00000016458 | Wt1 | Metanephric mesenchyme |
| Leydig | Cyp19A1 | ENSMUSG00000032274 | Cyp19a1 | |
| Leydig | StAR | ENSMUSG00000031574 | Star | |
| Leydig | Hsd17b3 | ENSMUSG00000033122 | Hsd17b3 | |
| Leydig | Hsd3b1 | ENSMUSG00000027871 | Hsd3b1 | |
| Leydig | Cyp17a1 | ENSMUSG00000003555 | Cyp17a1 | |
| Leydig | Lhcgr/ KH-R | ENSMUSG00000024107 | Lhcgr | |
| Leydig | Cyp11a1/P450Scc | ENSMUSG00000032323 | Cyp11a1 | |
| LPM | ISL1 | ENSMUSG00000042258 | Isl1 | |
| LPM | LMO2 | ENSMUSG00000032698 | Lmo2 | |
| LPM | Nkx2-5 | ENSMUSG00000015579 | Nkx2-5 | |
| LPM | PECAM1 | ENSMUSG00000020717 | Pecam1 | |
| Mesoderm | Cdx2 | ENSMUSG00000029646 | Cdx2 | Mesoderm +Paraxial Mesoderm |
| Mesoderm | Eomes | ENSMUSG00000032446 | Eomes | Mesoderm +endoderm |
| Mesoderm | Mesp1 | ENSMUSG00000030544 | Mesp1 | Mesoderm +Paraxial Mesoderm |
| Mesoderm | Mixl1 | ENSMUSG00000026497 | Mixl1 | |
| Mesoderm | Tbx6 | ENSMUSG00000030699 | Tbx6 | |
| Mesoderm | T/ Brachyury | ENSMUSG00000062327 | T | |
| PM | Cdx2 | ENSMUSG00000029646 | Cdx2 | |
| PM | Mesp1 | ENSMUSG00000030544 | Mesp1 | |
| PM | Tbx6 | ENSMUSG00000030699 | Tbx6 | |
| PM | TCF15 | ENSMUSG00000068079 | Tcf15 | |
| Sertoli | Aldh1a1 | ENSMUSG00000053279 | Aldh1a1 | |
| Sertoli | Amh | ENSMUSG00000035262 | Amh | |
| Sertoli | Cited1 | ENSMUSG00000051159 | Cited1 | |
| Sertoli | Clu | ENSMUSG00000022037 | Clu | |
| Sertoli | cst9 | ENSMUSG00000027445 | Cst9 | |
| Sertoli | Cyp26b1 | ENSMUSG00000063415 | Cyp26b1 | |
| Sertoli | Dhh | ENSMUSG00000023000 | Dhh | |
| Sertoli | Dmrt1 | ENSMUSG00000024837 | Dmrt1 | |
| Sertoli | Erbb4 | ENSMUSG00000062209 | Erbb4 | |
| Sertoli | FGF9 | ENSMUSG00000021974 | Fgf9 | |
| Sertoli | GATA1 | ENSMUSG00000031162 | Gata1 | Adult Sertoli |
| Sertoli | GATA4 | ENSMUSG00000021944 | Gata4 | |
| Sertoli | GDNF | ENSMUSG00000022144 | Gdnf | |
| Sertoli | Krt18 | ENSMUSG00000023043 | Krt18 | |
| Sertoli | Ptgds | ENSMUSG00000015090 | Ptgds | |
| Sertoli | SF1 | ENSMUSG00000024949 | Sf1 | |
| Sertoli | SHBG | ENSMUSG00000005202 | Shbg | |
| Sertoli | Sox8 | ENSMUSG00000024176 | Sox8 | |
| Sertoli | Sox9 | ENSMUSG00000000567 | Sox9 | |
| Sertoli | Vnn1 | ENSMUSG00000037440 | Vnn1 | |
| Sertoli | WT1 | ENSMUSG00000016458 | Wt1 | |
| Sertoli | Col9a1 | ENSMUSG00000026147 | Col9a1 | |
| Sertoli | Fgfr2 | ENSMUSG00000030849 | Fgfr2 | |
| Sertoli | Ncoa2/Tif2 | ENSMUSG00000005886 | Ncoa2 | |
